# Sex-specific behavioral feedback modulates sensorimotor processing and drives flexible social behavior

**DOI:** 10.1101/2025.05.08.652884

**Authors:** Sarath Ravindran Nair, Adrián Palacios Muñoz, Sage Martineau, Malak Nasr, Jan Clemens

## Abstract

How the brain enables individuals to adapt behavior to their partner is key to understanding social exchange. For example, courtship behavior involves sensorimotor processing of signals that can result in behavioral dialogue between partners, such as stereotyped movements and singing. The courtship behavior of *Drosophila melanogaster* males with their partners, which are usually female but can also be male, involves singing. To investigate how behavioral feed-back and sensorimotor processing contribute to flexible social interactions, we compared the courtship behavior and singing of male *D. melanogaster* towards males and females. Quantitative analysis of their interactions revealed that while underlying courtship and song rules are unaffected by the sex of the partner, the behavioral dynamics and song sequences differ by partner sex. This divergence stems from sex-specific behavioral feedback: females decelerate to song, while males orient towards the singer. Moreover, optogenetic manipulations reveal that the partners’ responses are driven by sex-specific neural circuits that link song detection with arousal and social decisions. Our findings demonstrate that flexible social behaviors can arise from fixed sensorimotor rules through a context-dependent selection facilitated by the partner’s behavioral feedback. More broadly, our results reveal compositionality as a key mechanism for achieving behavioral flexibility during complex social interactions such as courtship.

## Introduction

The brain enables flexible behavior by dynamically transforming sensory cues into motor outputs through sensorimotor rules [1–5]. These rules are not necessarily fixed [6–10], but flexibly selected and modulated based on the organism’s internal state and the external sensory context [11–16]. In social settings, each individual’s behavior generates sensory cues for others, creating a closed loop of reciprocal influence. To navigate dynamic social environments, the brain must interpret and flexibly respond to social signals in real time. This flexibility can be achieved by applying different rules for different partners—based on innate preferences [17–20], learning [21–23], or prior experience [24–26]—and by modulating behavior in response to varying social feedback when applying each rule [13, 27]. Understanding the mechanisms underlying flexible rule selection and application is key to unraveling how the brain controls flexible behavior.

Courtship interactions are an example of a behavior shaped by behavioral feedback and sensorimotor processing. Although typically directed at members of the opposite sex, same-sex courtship is also widespread in the animal kingdom (see Sommer and Vasey [28] for a comprehensive review). Homosexual courtship is often assumed to be identical to heterosexual courtship behavior, but the extent to which individuals employ the same rules for homosexual versus heterosexual courtship is unclear. Importantly, members of the same sex often respond differently to being courted than members of the opposite sex [29–31], yet whether this differential feedback shapes the behavior of the courter remains poorly understood. Thus, comparing courtship behaviors directed at different sexes offers a compelling entry point for understanding how the brain produces flexible behavior through the interplay of behavioral rules and feedback [1, 5, 32–34].

In *Drosophila melanogaster*, chemical cues indicate the sex and species of an individual [35– 37]. Males sample the chemical profile of other flies and typically initiate courtship toward females of their own species [36, 38, 39]. The male chases the female and produces a courtship song by extending and vibrating one wing. This courtship song contains two main modes [40]: pulse song, consisting of regular trains of two types of short pulses [41], and sine song, a sustained oscillation of the wing. The sensorimotor rules that transform feedback from females to song patterning in the courter determine the timing, duration, and composition of the song [4, 13, 41–43]. This involves males switching between a set of three rules, *close, chasing*, and *whatever*, depending on the male’s internal state and interaction context [13]. Males use the *whatever* rule when not interested in an interaction, the *chasing* rule when the female is fast and distant to produce mainly pulse song, and the *close* rule when she is slow and nearby, producing mainly sine song.

Male-male interactions in *Drosophila* in the presence of a resource—food or a female—are often aggressive and involve head butting, boxing, fencing, wing threats, lunging, and chasing [44–46]. During aggressive interactions, males produce agonistic song, which is more irregular than courtship song and generated by bilateral wing flicking [44, 47–49]. However, males also frequently court other males in the wild [50]. In laboratory settings, courtship-like behavior between males can be induced via genetic mutations but can also occur spontaneously. In groups, song playback leads to courtship-like chaining behavior [25, 51–53], during which males chase each other’s tails and extend one wing. However, whether the unilateral wing extensions during male-male interactions produce song, and if so, whether male-directed song comprises the same modes and patterns as female-directed song, has not been systematically examined.

By comparing male- and female-directed courtship and modes of singing, we elucidate how social feedback and internal sensorimotor rules interact to generate flexible social behavior. We find that male- and female-directed courtship unfolds with different dynamics. This results in differences in song patterns, which arise because of the partners’ sex-specific behavior: Female partners typically stop during courtship, while male partners often turn back to face the courting male. This turning behavior in males leads to different sensory experiences from singing to females, which changes when specific rules are used and thus singing behavior. Moreover, we identify a putative neural circuit that links song perception with different behavioral responses in males. Taken together, our results demonstrate how flexible behavior emerges from compositionality, such that fixed sensorimotor rules are used according to social feedback. Our study thus proposes a mechanism through which flexibility can be achieved even in innate behaviors such as courtship.

## Results

### Male- and female-directed courtship-like interactions exhibit different dynamics

To compare male- and female-directed singing, we tracked the interactions of male-male and male-female pairs [54] and recorded acoustic signals with an array of 16 microphones [4, 55, 56]. To promote male-male interactions, we controlled the males’ prior social experience and perfumed the chamber with male and female flies prior to the experiments (see Methods). In our assay, males interacted intensely with both males and females (females 86±12% vs. males 73 ± 15%, Fig. 1A). Female-directed interactions had lower latency and were more frequent, indicating that the males still discriminated between sexes (Fig. 1A). Using uni- and bilateral wing extension as proxies of courtship and aggression, we found that even the male-male interactions were courtship-like: Males primarily displayed unilateral wing extensions, with bilateral extensions being nearly absent in male-female pairs and rare in male-male pairs (Fig. 1B). The audio recordings confirmed our interpretations of the wing extensions: Males produced mainly courtship song when interacting with either sex, while agonistic song was absent during male-female interactions and rare during male-male interactions (Fig. 1B).

**Figure 1:**
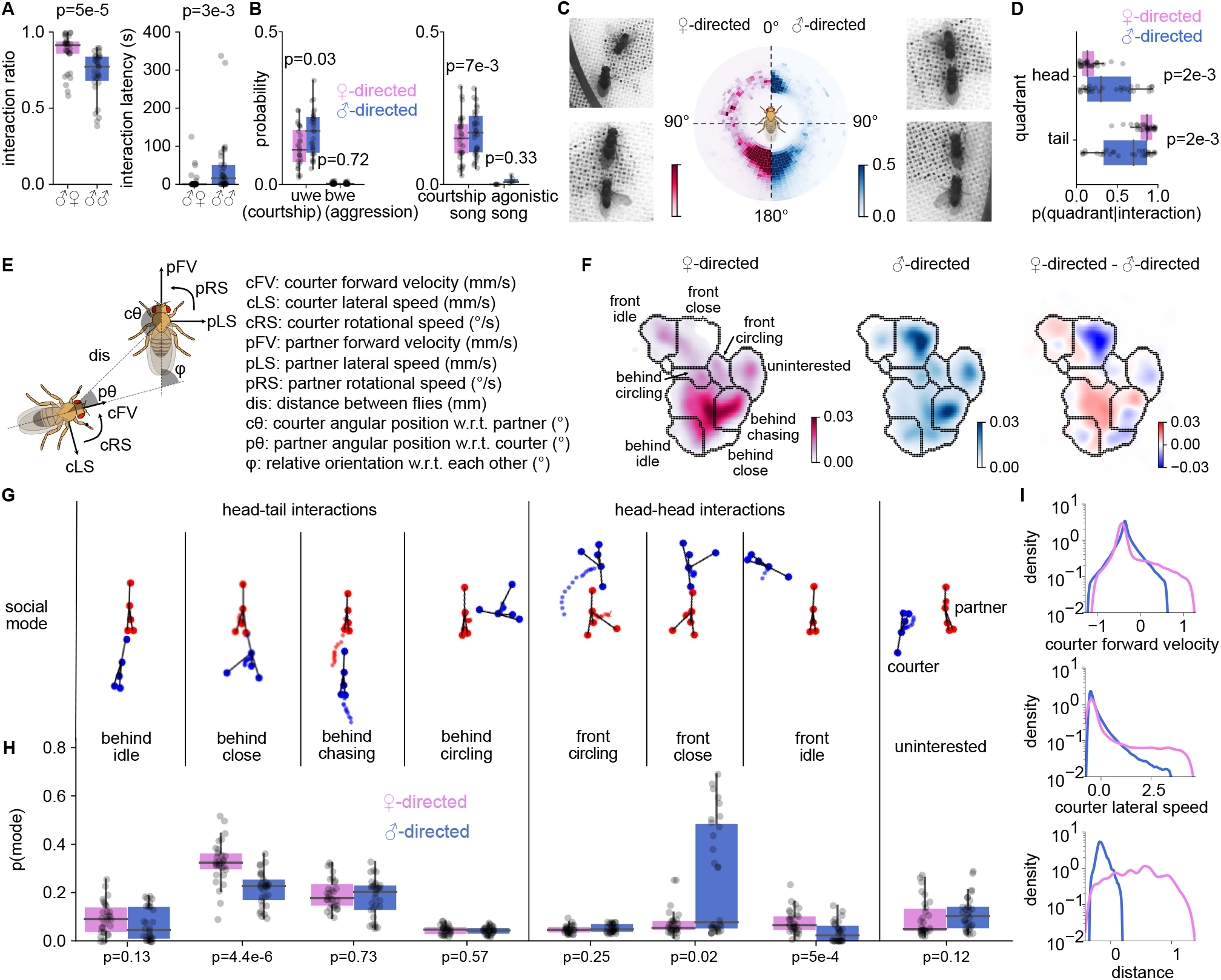
Male- and female-directed courtship-like interactions exhibit different dynamics. **A** Fraction of time spent interacting (left) and latency to interaction (right) for male-female (left, purple) and male-female pairs (right, blue). **B** Wing movements (left) and acoustic signals (right) produced in male-female (magenta) and male-male (blue) pairs. Unilateral wing extensions (uwes, left) indicate courtship-like interactions and typically lead to the production of courtship song (right). Bilateral wing extensions (bwes) indicate aggression and typically lead to agonistic song. **C** Position of a male courter around a female (left, magenta) or a male (right, blue) partner (see colorbar). 0° and 180° represent the partner’s head and tail, respectively. Males are typically behind the female partner (tail interaction) or circle in front (arc) of her. By contrast, males spend more time directly in front of a male partner (head interaction). Males also tend to be positioned slightly farther from a male vs a female partner during tail interactions. Images show examples of tail (bottom) and head (top) interactions in male-female (left) and male-male (right) pairs **D** Ratio of time spent by the male courter near the head and tail of the partner during interactions. **E** Parameters used to generate the social maps in I (see Methods for details). **F** Social maps for female-directed interactions (left), male-directed interactions (middle), and their difference (right). Color codes densities (left, middle) or their differences (right), see color bars. **G** Social modes represented by each cluster in the interaction state space. **H** The fraction of time spent in each social model during female-(magenta) and male-directed (blue) interactions. **I** Distribution of courter forward velocity, courter lateral speed, and distance when the male courter is singing near the head of the partner. The male is faster and at larger distances when singing near the head of a female partner and is slower and closer when singing near the head of a male partner. p-values were obtained using two-sided Mann-Whitney U tests. Dots in A, B, D, and H correspond to average values for each pair of flies (N=30 male-male and 30 male-female pairs).

However, although male- and female-directed interactions were both courtship-like, they unfolded with different dynamics. During male-female interactions, the male spent 84±12% of the time behind the female, facing her tail. By contrast, male-male pairs frequently (42 ± 31%) engaged in head interactions, with both males facing each other (Fig. 1C, D, S1). These head interactions were not specific to the NM91 wild-type strain we used in the assay: OregonR wild-type males also engaged in frequent head-directed interactions with male but not female partners (males: 14±10% vs females: 7±5%, Fig. S1). Although head interactions between males typically indicate aggression [46, 57], males produced primarily courtship song during the head interactions (Fig. S3A), suggesting that these encounters were courtship-like rather than aggressive.

To more comprehensively compare the dynamics of male- and female-directed interactions, we generated social maps from the joint egocentric and relational kinematics of both partners using the UMAP transform (Fig. 1E—F, S2) [58–61]. We then grouped similar social behaviors into eight social modes (Fig. 1F—G, S2), which revealed similarities and differences in the interactions with male and female partners (Fig. 1F, H): Males spent similar time chasing male or female partners (“behind chasing”). However, during male-female interactions, males were more often sitting “behind close” or in front of the female (“front idle”) [62]. By contrast, males were more often oriented towards each other and close (“front close”). Males engaged in head interactions with both males and females, and the head interactions differed not only in frequency (Fig. 1H) but also in their dynamics (Fig. 1I). During head interactions with a female, males were more distant and either idle or circled around a stationary female. During male-directed head interactions, males were closer and moved more slowly (Fig. 1I). Therefore, the dynamics of male- and female-directed courtship-like interactions are different, thus creating distinct sensory experiences for a courting male.

### Song patterns depend on the partner’s sex during head interactions

Given that courtship song is shaped by the sensory experiences of the male [4, 13, 41–43], we investigated whether male- and female-directed song patterns have distinct properties, particularly during head interactions. Song is patterned on two timescales: On a short timescale (<50 ms), the frequency content of the sine song and pulses as well as the interval between pulses are produced by largely hard-wired circuits in the ventral nerve cord [63]. Accordingly, male- and female-directed songs were nearly identical on the short timescale, with similar carrier frequencies of pulse and sine, pulse durations, intervals, and waveform shapes (Fig. 2A–B, S3).

**Figure 2:**
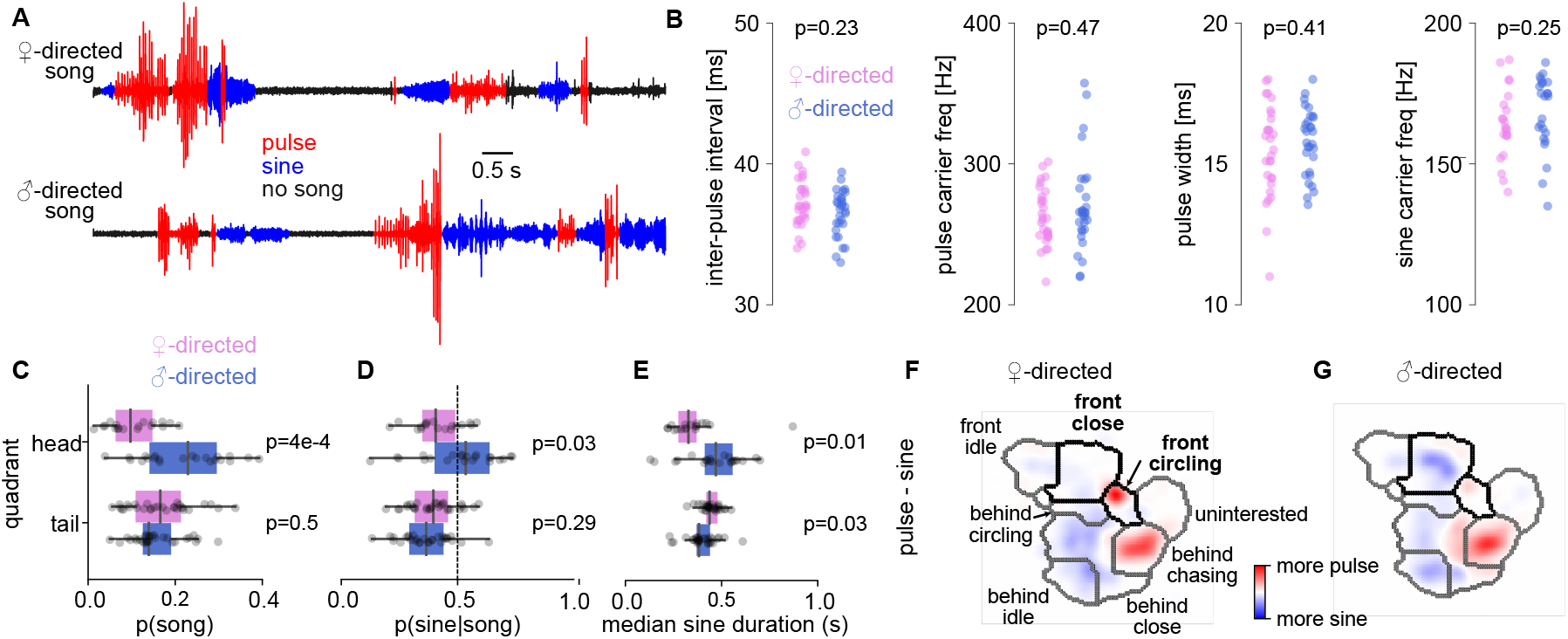
Song patterns depend on the partner’s sex during head interactions. **A** Traces of female (top) and male (bottom) directed courtship song. Both female- and male-directed song contain the pulse (red) and sine (blue) modes. **B** Song features on the short timescale (<50 ms) are identical in male (blue) and female-directed (blue) courtship song. **C, D** Probability of courtship song (C) and fraction of sine song out of all song (D) during head (top) and tail (bottom) interactions with a male (blue) and female (magenta) partner. Dashed line in D indicates an equal amount of pulse and sine. **E** Median duration of sine songs during head (top) and tail (bottom) interactions with a male (blue) and female (magenta) partner. **F, G** Difference between the social maps for pulse and sine song during female-(**F**) and male-directed (**G**) interactions showing sex-specific differences in song choice across social modes (see color bar). p-values were obtained using two-sided Mann-Whitney U tests. Dots in B–E correspond to average values for each NM91 fly pair. Each pixel in the social maps in F–H is averaged across fly pairs. N=30 male-male and 30 male-female pairs.

On a longer timescale (>50 ms), sine and pulse are sequenced into song bouts. We detected differences in song sequences only during head interactions (Fig. 2C–E, S4). During head interactions, males spent more time singing to a male than to a female partner (female: 10 ± 6% vs male: 22 ± 10%, Fig. 2C) and they sang more and longer sine song to a male than to a female (Fig. 2D, E). Additionally, males sang in different social modes near the head of a female and male partner (Fig. 2F, G, S5): Pulse-biased female-directed song occurred mainly during the “front circling” mode. Sine-biased male-directed song occurred mainly during the “front close” mode.

### Males use the same sensorimotor rules for male and female-directed singing

The sex-specific song patterning could arise from the males using sex-specific sensorimotor rules or from male and female partners providing differential feedback. To discriminate between these hypotheses, we first identified the song-patterning rules by combining a hidden Markov model (HMM) with generalized linear models (GLMs) [13, 64–66] (Fig. 3A). In an HMM-GLM, each song patterning rule is represented by a GLM, which transforms partner feedback into the courting male’s singing behavior: whether to produce pulses, sine songs, or nothing (Fig. 3B). The model’s HMM component determines which of several rules is used during each time point. This combination of an HMM with GLMs can account for possible sex- and context-specific behavioral rules by allowing the model to flexibly switch cue-to-song mappings (Fig. 3C) according to the sex and behavior of the partner. As surrogate feedback cues, we used the kinematics of the courting male and the partner fly as well as their relative positions (Fig. 3B, 1E) [4, 13].

**Figure 3:**
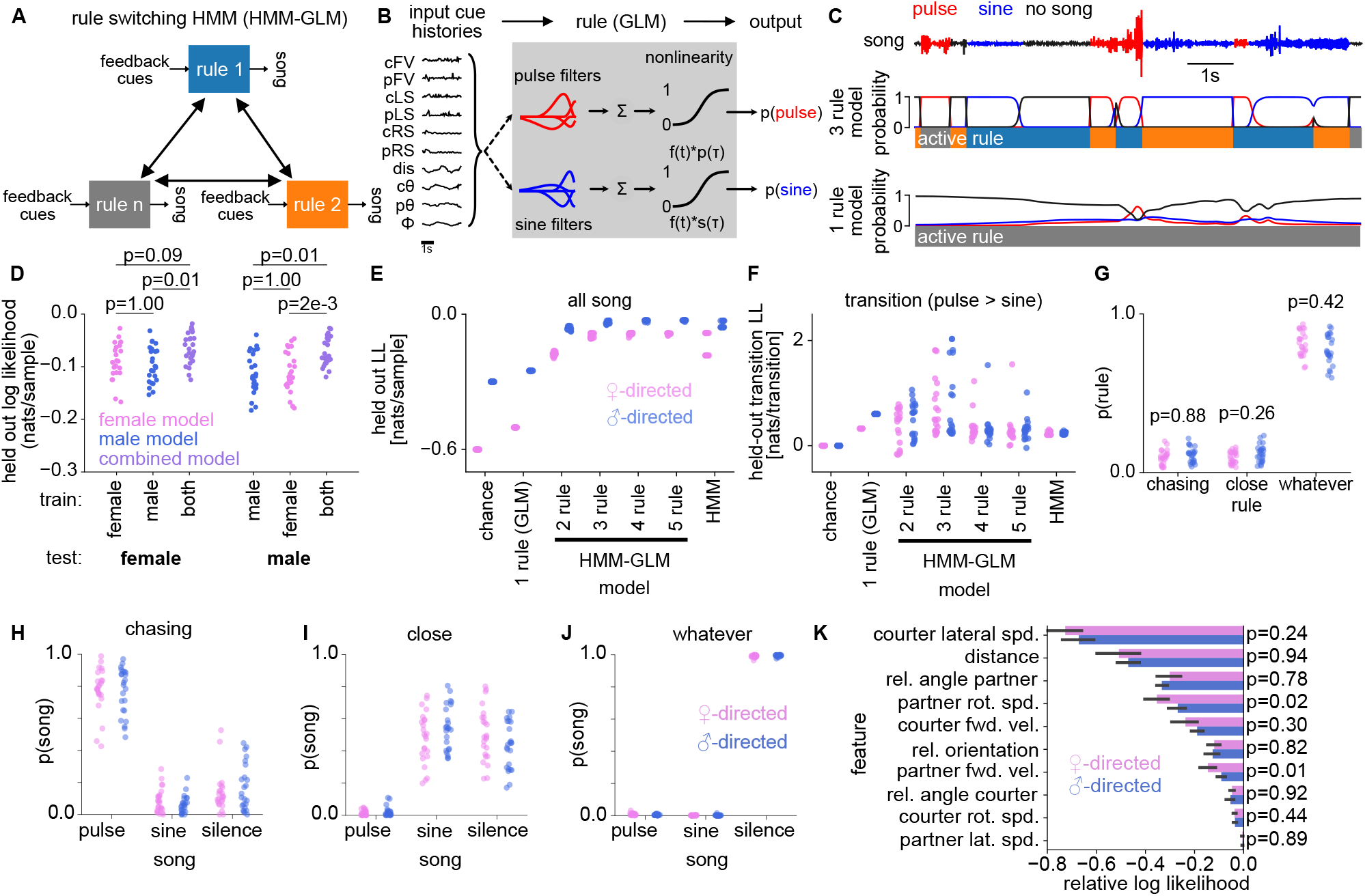
Males use the same sensorimotor rules for male and female-directed singing. **A** Male- and female-directed singing were modeled using an HMM-GLM, a combination of a hidden-Markov model (HMM) and a generalized-linear model (GLM). Males switch between a fixed set of rules (boxes), which define sensorimotor transformations that map feedback cues to song. **B** Each rule is implemented as a multinomial GLM, which predicts the song pattern (pulse, sine, or no song) from the dynamical feedback cues given by the interaction features extracted from the tracking data (Fig. 1E). Each feature is filtered through a set of linear filters (one for pulse and one for sine song). A non-linear function determines the probability of each song mode based on the sum of the filtered cues. No song is given by 1 minus the probability of pulse and sine. **C** HMM-GLM with one (bottom) and three (middle) rules fitted to male- and female-directed song data. Shown are 10 seconds of interaction during which the male switched between different song modes (pulse - red, sine - blue, no song - black) (top). The three-rule HMM-GLM faithfully predicts the male song pattern. **D** Goodness of fit (log likelihoods) for a three-rule HMM-GLM trained using male-directed data (blue), female-directed data (magenta), or a combination of both (purple). Dots show likelihoods calculated on held-out data from female (left) or male-directed (right) interactions (N=25 held-out trials each). All models predict song patterns for both types of held-out test data, irrespective of the training data used. Models trained with combined data predicted the song patterns slightly better compared to sex-specific models, likely due to the larger size and diversity of training data. **E, F** Normalized log-likelihood of the models on held-out data of female- and male-directed singing for all song (E) and only during transitions between song modes (F, pulse to sine transition shown here). Dots correspond to fits with different initializations (N=20). The state-aware HMM-GLM models could better explain both the female- and male-directed song sequencing compared to a GLM or HMM model. An HMM-GLM model with more than three states could not explain the song sequencing better than those with three states. **G** The fraction of time spent employing different rules during female- and male-directed interactions. Dots correspond to pairs (N=30) **H–J** Probability of singing each song mode when using the chasing (**H**), whatever (**I**), and close (**J**) rule towards males (blue) and females (magenta). The chasing rule produces predominantly pulse, but also sine and silence, the close rule predominantly sine and silence, and low amounts of pulse song as well, and whatever rule predominantly silence. Dots correspond to pairs (N=30 male-female and 30 male-male pairs). **K** Importance of each feature for predicting the song mode, computed as the reduction in log-likelihood of the model when the given feature is randomly permuted. All p-values were obtained using two-sided Mann-Whitney U tests

When fitting the HMM-GLM to predict singing in our behavioral data, we found that males use the same set of rules for male- and female-directed singing: A model trained with combined data from male- and female-directed singing predicted song patterns as well as sex-specific models trained to explain singing towards only one of the two sexes. In addition, the sex-specific models generalized to explain singing to the sex not used for training (Fig. 3D). An HMM-GLM predicted song patterns better than an HMM or a GLM alone (Fig. 3C, E–F), implying that both the ability to switch rules (HMM) as well as the ability to flexibly respond to feedback cues (GLM) are crucial for explaining male singing. Models with three rules predicted the patterns of male- and female-directed song as accurately as a model with more rules (Fig. 3E). Additionally, the model with three rules predicted transitions between song modes better than models with fewer or more rules (Fig. 3F). The rules identified for male- and female-directed singing map to the rules previously identified for female-directed singing [13] (Fig. 3H-J, S6A): A non-interaction *whatever* rule, a pulse-biased *chasing* rule that is used when males are faster and the partner is farther, and a sine-biased *close* rule that is used when the male is slower and the partner is closer. Males use these rules with equal frequency towards either sex (Fig. 3G).

The same feedback cues determine singing for both partner sexes (Fig. 3K, S6B): The courting male’s lateral speed and distance from the partner are the most important features for predicting singing towards male and female partners. By contrast, the cue that was most dependent on partner sex—the male’s position around the partner (head or tail, Fig. S6C)—is less important for song patterning. This indicates that the differences in song patterns do not arise simply from differences in the male’s position around females and males, but from differences in his speed and distance from the partner during head interactions. Given that males use identical rules and cues for singing towards male and female partners, sex-specific song patterns likely arise from differences in partner feedback. We therefore examined and manipulated partner feedback.

### Partner feedback drives sex-specific song patterns

To characterize sex-specific partner responses to song, we constructed behavioral maps [60, 61, 67] of the partner’s behavior during male singing (Fig. 4A). This revealed profound sex-specificity in the responses to courtship song: Female partners spend more time being idle and moving slowly or remaining stationary than male partners (Fig. 4B, S7). The female’s idleness enables the male to initiate head interactions by rapidly circling towards her head while maintaining sufficient distance, likely to avoid startling her (Fig. 4C, D, S7C–D, [62, 68]). Since these head interactions are faster and occur at a distance, males use the chasing rule (Fig. 4E) and predominantly sing pulse song (Fig. 3H). By contrast, male partners typically run away or frequently sing back by extending their wing (Fig. 4A–B, S7). The male partner’s constant movement prevents the singing male from circling in front. Instead, male-directed head interactions are initiated by the partner turning back to face and approach the singer (Fig. 4C, D). This slows the courting male and reduces the distance between both males, forcing the courter to choose the close rule (Fig. 4E) and sing more sine song (Fig. 3I). These observations suggest that partner feedback drives differences in rule use which in turn leads to sex-specific song patterns (Fig. 4F). Moreover, it implies that the head interactions between male partners and the subsequent increase in sine song are enforced by the male partner, not from deliberate choices by the courter.

**Figure 4:**
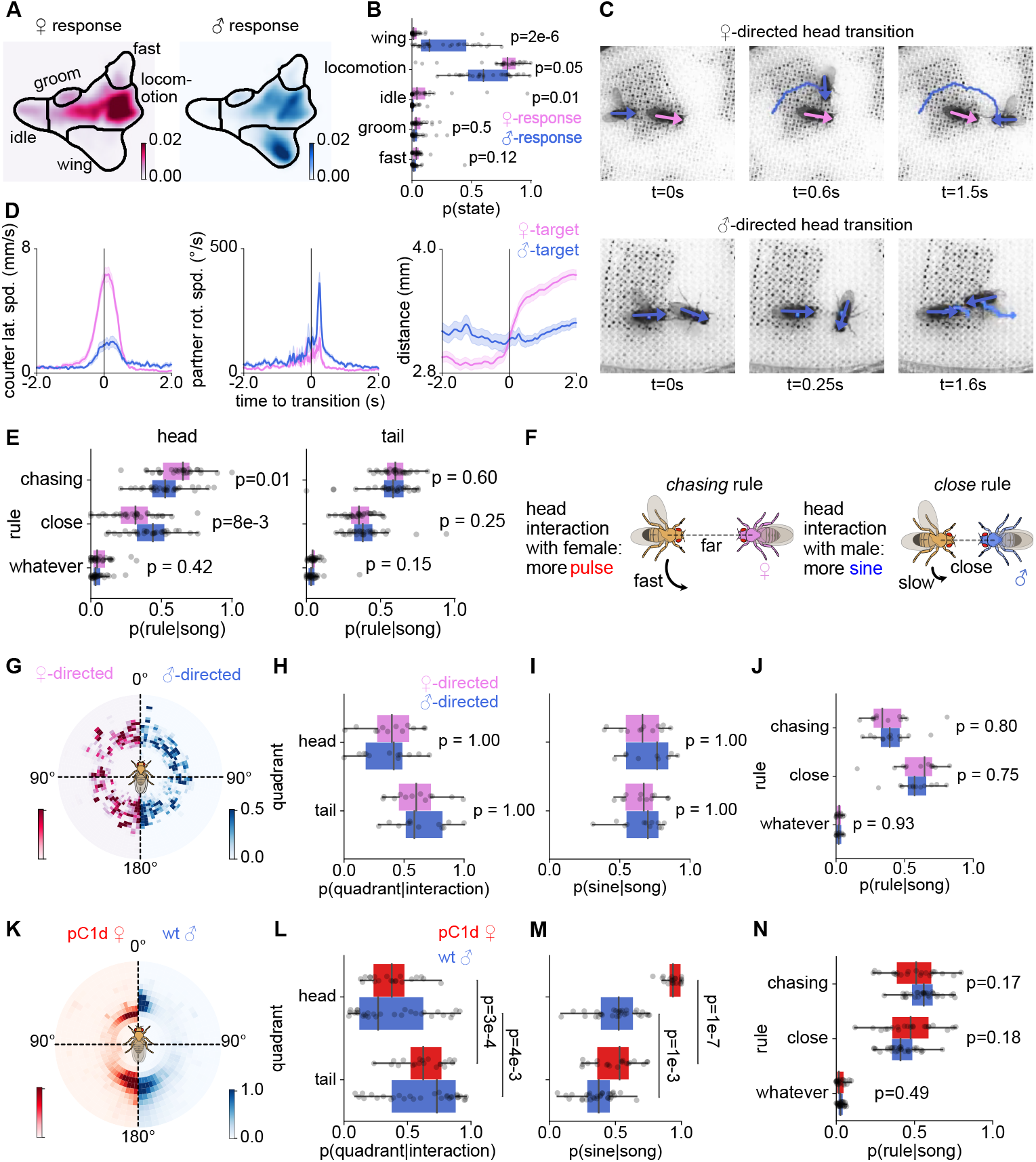
Partner feedback drives sex-specific song patterns. **A–B** Probability of different behaviors of female (right) and male (left) partner while the courter male is singing. Females spent more time in an idle state, and males spent more time extending their wing as a response to the song. Dots in **B** correspond to each fly pair (N=30 male-female and 30 male-male pairs). **C** Representative transitions into head interactions towards a female (top) and a male (bottom) partner. Transitions into head interactions depend on the partner’s sex. Head interactions with a female are initiated by the male moving to the front of the stationary female partner, whereas those between males are initiated by the male partner turning back. **D** Courter lateral speed (left), partner rotational speed (middle), and distance (right) during transitions into head interactions with male (blue) and female (magenta) partners show sex-specific dynamics. Transition to head interactions with a female is accompanied by an increased courter lateral speed, and an increase in distance as the male circles to the front of the female. Transition to head interactions with a male is accompanied by an increased partner rotational speed as the male partner turns back. Lines and shaded areas correspond to mean ±s.e.m. (standard error of mean) (N=94 transitions from 25 male-female pairs and N=88 transitions from 26 male-male pairs). **E** Fraction of rule use when the courter is singing near the head (left) and tail (right) of a male (blue) or female (magenta) partner. Dots correspond to pairs (N=30 male-female and 30 male-male pairs) **F** Sex-specific song sequences arise from differential rule use according to partner feedback. When singing near the head of a female partner (left), the male is fast and further away from the female (as the male courter circles in front of the female partner), and mainly produces pulse song. When singing near the head of a male (right), the male courter is slower and closer (as the male partner turns back and approaches the courter), and mainly produces sine song. **G** Position of a male courter around an immobilized female (magenta) or male (blue) partner. **H** Fraction of time spent by the courter male in the head and tail quadrants around an immobilized female (magenta) and male partner (blue) during interactions. **I** Fraction of sine song (out of all song) sung by the courter male in the head and tail quadrants around an immobilized female (magenta) and male partner (blue). **J** Rules use when singing near an immobilized partner. Dots in H–J correspond to each fly pair (N=12 each for male-female and male-male pairs). **K** Position of a male courter around a pC1d-activated female compared to that around a wild-type male. **L** Fraction of time spent by the courter male in the head and tail quadrants around a pC1d-activated female (red) and a wild-type male partner (blue) during interactions. **M** Fraction of sine song (out of all song) sung by the courter male in the head and tail quadrants around a pC1d-activated female (red) and a wild-type male partner (blue). **N** Comparison of rule use when singing to a pC1d-activated female partner and a wildtype male partner. Dots in L–N correspond to each fly pair (N=20 for pC1d-activated females and N=30 for wild-type males). p-values were obtained using two-sided Mann-Whitney U tests, except for panels L–M, for which a two-sided Wilcoxon signed rank test was used.

To determine whether the causal factor is behavioral feedback rather than the partner’s sex [68, 69], we manipulated partner behavior by slowing male partners to resemble females, or making female partners turn back like males. We inactivated motor neurons to slow and stop one male in a pair using the light-gated anion channel GtACR1 [70]. Inducing idleness in a male partner was sufficient to evoke frontal circling by the non-manipulated male (Fig. 4G–H, S8A) and eliminated sex-specific differences in rule use and song sequences (Fig. 4I–J). Next, we induced female turning by optogenetically activating pC1d neurons using the light-gated cation channel CsChrimson [71], which induces aggression and turning in females [72, 73]. During pC1d activation, females turned back toward the courting males, leading to head interactions similar to those seen with male partners (Fig. 4K–L, S8B–C). The manipulated female partners were even closer to the courter than male partners, leading males to sing even more sine song using the close rule (Fig. 4M–N, S8D–E). Thus, sex-specific differences in song patterning are driven by behavioral feedback from the partner, not by their sex or by the courters’ choice.

### Courtship song perception drives male-male head interactions

To test whether courtship song drives the turning of the male partner, we manipulated song perception in male pairs. In pairs with one aristacut (deaf) or one wingcut (mute) male, head interactions were extremely rare (Fig. 5A, E). The few head interactions detected were mostly initiated by the male in the pair who could still hear the song produced by his partner (Fig. 5D, H). The male in the pair that did not hear the song (the deaf male paired with intact or the intact male paired with mute male) did not initiate head interactions but typically fled from the other male, leading to tail interactions (Fig. 5B–C, F–G). Head interactions were also rare in pairs of two mute males but could be induced by pulse song playback that resulted in turning responses of the partner (Fig. 5I–L). Together, these experiments demonstrate that perceiving song is both necessary and sufficient for head interactions.

**Figure 5:**
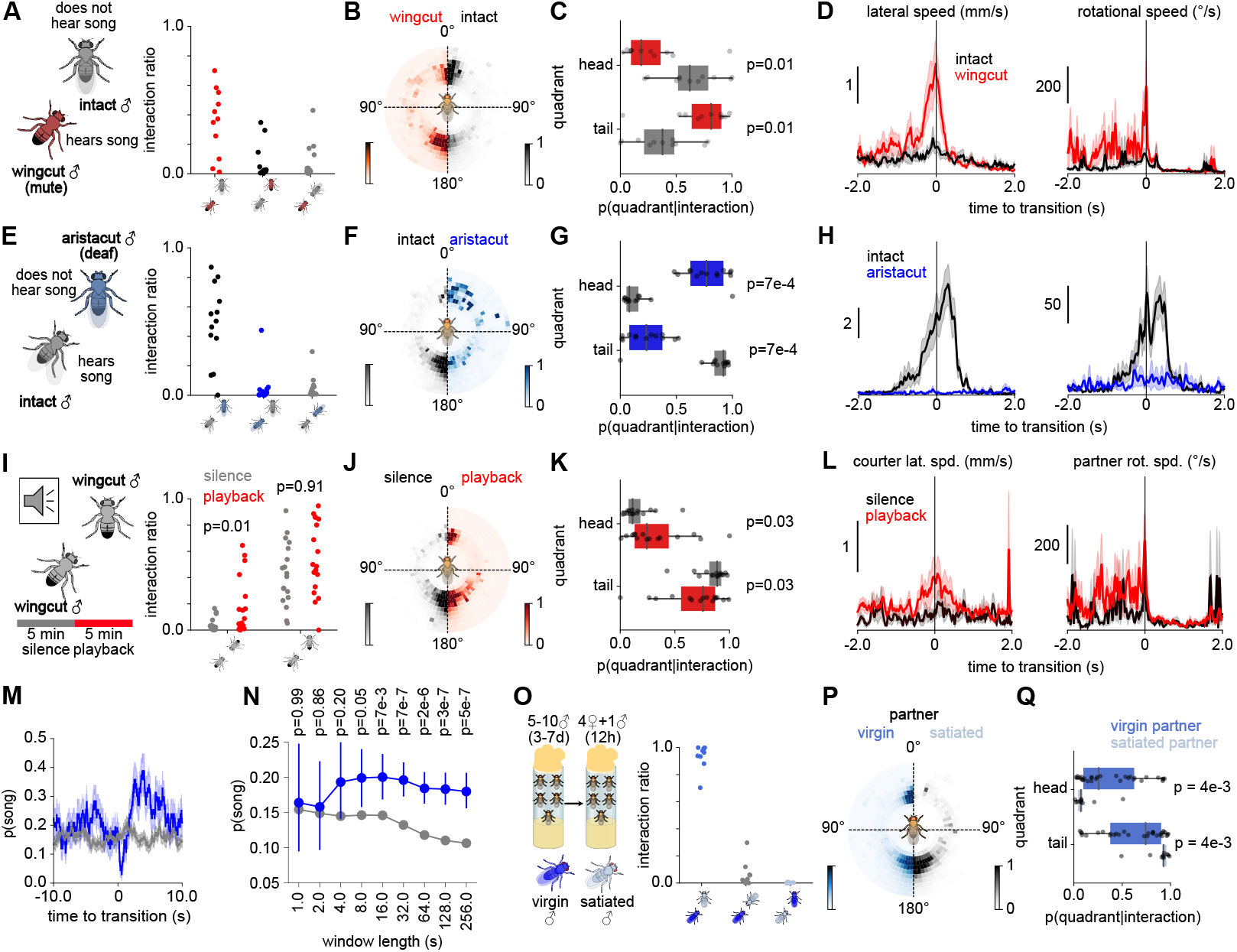
Courtship song perception drives male-male head interactions. **A** Fraction of trial time spent in different configurations during interactions between an intact (grey) and a wingcut (mute) male (red) (from left to right: mute male behind intact male, intact behind mute, and mute and intact facing each other). **B** Position of the mute male when courting the intact male (red, left) and the position of the intact male when courting the mute male (black, right). **C** Fraction of interaction time spent by the wing-cut and the intact male in the head and tail quadrant during courtship. The mute male is placed mostly near the tail when courting the intact partner male, whereas the intact male is placed mostly near the head when courting the mute partner (N=11 pairs). **D** Lateral (left) and rotational speed (right) of the mute (red) and intact male (black) during transitions to head interactions. Head interactions are mostly initiated by the mute male (N=24), either by circling to the front of the intact or by turning back to the intact male. Intact flies rarely initiate head-to-head interactions when courted by the mute male (N=12). **E** Same as A, but for interactions between an intact (grey) and an aristacut (deaf) male (blue). **F** (Left) Position of the intact male when courting the deaf male (black, left) and vice versa (blue, right). **G** Fraction of interaction time spent by the intact and deaf male in the head and tail quadrant during courtship. The intact male is placed mostly near the tail of the deaf male, the deaf male is placed mostly near the head of the intact male (N=13 pairs). **H** Lateral (left) and rotational speed (right) of intact (black) and deaf (blue) males during transitions to the head interactions. Head interactions are initiated by the intact male either by circling to the front of the deaf male or by turning back to it (N=9). Deaf males rarely initiate head-to-head interactions (N=2). **I** (Left) Playback experiments with pairs of wingcut (mute) males. Males interacted for five minutes in silence (silence control) after which a conspecific pulse song was played back continuously for five minutes (playback experiment). (Right) Fraction of trial time spent in different configurations (both flies facing each other and one fly behind the other) during silence (black) and playback (red). **J** (Left) The position of the courter male during silence (black, left) and song playback (red, right). **K** The fraction of interaction time spent by the courter in the head and tail quadrant during silence (black) and playback (red). The courter spends significantly more time near the head of the partner during playback (N=15 pairs) **L** Courter lateral speed (left) and partner rotational speed (right) during transitions to head interactions. Head interactions are initiated mostly during playback by the courter moving laterally to the front or by the partner turning back to the courter (silence: N=6 transitions, playback: N=13 transitions). **M** The probability of courter singing in a window around the transition to head interactions. The grey line shows the song probability in randomly selected windows during which the courter maintained tail interactions (head transitions: N=64 windows from 22 male-male pairs, no transitions: N=2833 windows from 30 male-male pairs, all wild type). **N** The amount of song (mean *±* std) in windows of different durations preceding transitions to head interactions. The grey line shows the song amount in randomly selected windows where a head-to-head transition did not occur. Song was significantly enriched before the transitions only when evaluating longer windows, suggesting that song was integrated over longer time scales. **O** (Left) To assess the impact of sexual satiation on head interactions, males were isolated after eclosion and group-housed for 3-7 days. Sexual satiation was induced by housing one male with four virgin females for at least 12 hours before the experiment. (Right) Fraction of trial time spent in different configurations (from left to right: virgin male behind satiated, virgin and satiated males facing each other, and satiated male behind the virgin male) (N=10 pairs) **P** (Left) Position of the courter male when courting a naive (blue, left) and sexually satiated (black, right) male. **Q** The proportion of interaction time spent by the courter in the head and tail quadrant when courting a mature virgin (blue) and satiated male (grey). The courter is placed mostly near the tail of the satiated male partner and rarely near the head (virgin partner: N=30 pairs, satiated partner: N=10 pairs). For panels C, G, N, and Q, p-values were computed using two-sided Mann-Whitney-U tests, for panel I and K, using a two-sided Wilcoxon signed-rank test. Lines and shaded areas show the mean ±s.e.m. across all transitions. Dots correspond to fly pairs.

On what timescales does the song affect turning? In females, song affects locomotion on two timescales: Immediately, as a trigger for slowing within milliseconds [31, 74], and via integration over tens of seconds [75, 76]. In males, only direct, immediate effects of song on locomotor behavior [31], aggression [45], or chaining [25, 77] have been reported. Contrary to our expectation, song was not enriched immediately preceding male transitions from tail to head interactions (Fig. 5M). However, when evaluating song over longer timescales, we found more song in windows leading up to these transitions (Fig. 5N), indicating that turning was facilitated by song information integrated over tens of seconds. Thus, the mechanism underlying transitions from tail to head interactions is not simply phonotaxis [74] but likely involves a slow, song-driven buildup of the partner’s social arousal. This is consistent with head interactions being rare in sexually satiated males with low sexual arousal levels (Fig. 5O–Q).

### Neural circuitry that links acoustic information with arousal and courtship induces turning

Central neurons that detect courtship song in males are well characterized: The auditory vPN1 and pC2l neurons detect pulse song and induce courtship behaviors [6, 31, 52] (Fig. 6A). Both neurons are thought to activate different subsets of pC1 neurons, which act as social command neurons that drive courtship or aggression [52, 78–80]. To investigate the neural basis of song-induced male turning behavior, we tested the role of auditory vPN1 and pC2l neurons in driving head interactions and investigated which subsets of pC1 neurons drive these interactions.

**Figure 6:**
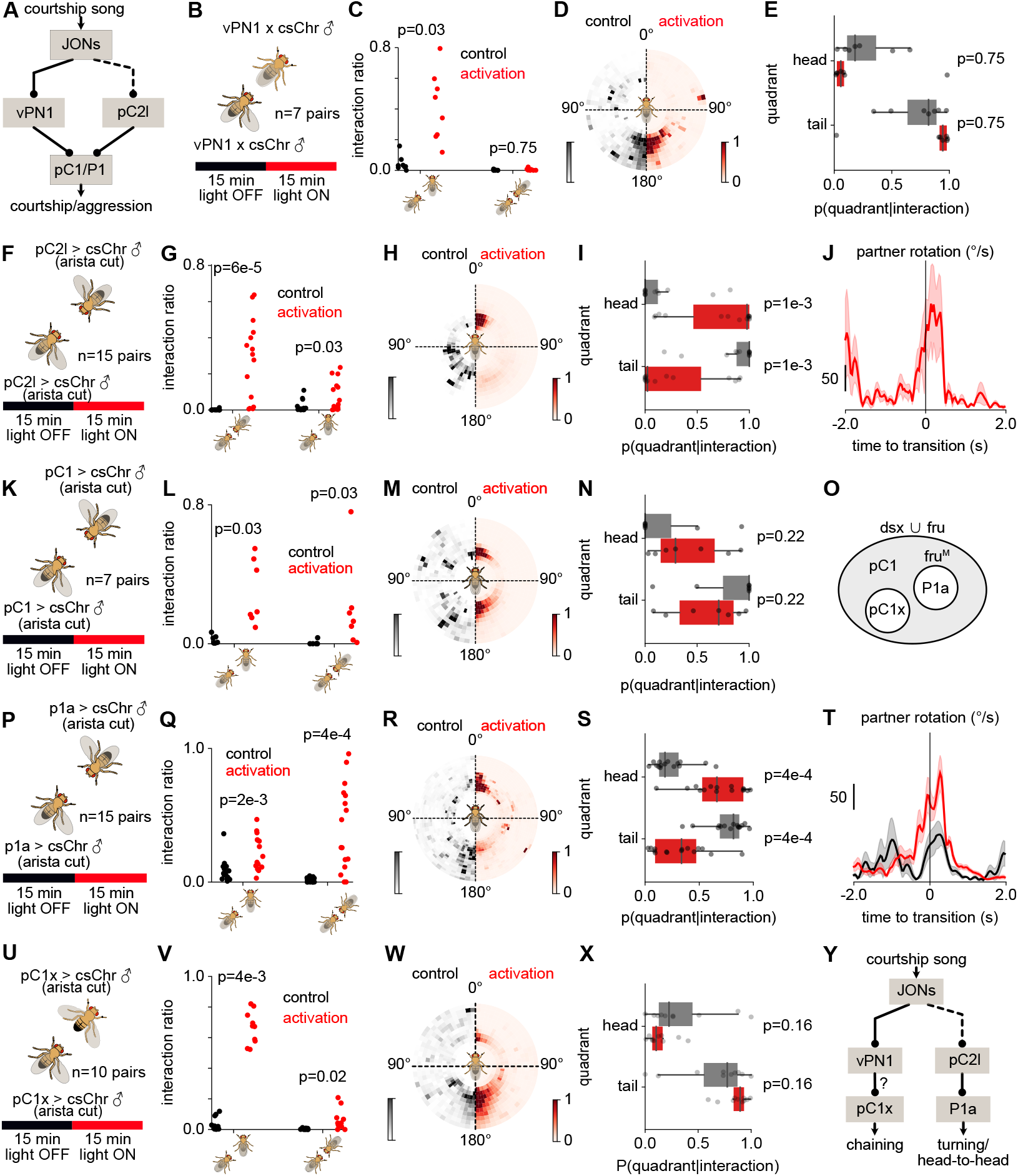
Song induces head interactions through a specific neural pathway. **A** Two parallel song detection pathways in the fly brain through male-specific vPN1 neurons and sexually dimorphic pulse song detector neurons pC2l [31, 52]. The vPN1 neurons induces male-male chaining behaviors [52] whereas pC2l induces sex-specific behavioral song responses such as slowing down in females and unilateral wing extension in males [31]. **B** Experimental procedure for optogenetic activation of vPN1 neurons (N=7 pairs). Two males that expressed red-shifted channel rhodopsin csChrimson in vPN1 neurons are paired for 30 minutes in the assay. For the first 15 minutes of the trial, the red LED was OFF (control) followed by vPN1 activation for 15 minutes by turning ON the red LED. **C** Fraction of trial time spent in head (left) and tail interactions (right) during control (black) and vPN1 activation (red). **D** Position of the courter male around the partner during control and activation of vPN1 neurons. **E** Fraction of interaction time spent in head (top) and tail (bottom) quadrants during control (black) and vPN1 activation (red). Dots in C and E correspond to averages for each fly pair (N=7 pairs). **F–I** Same as B–E, but for activation of pC2l neurons. Dots in G and I correspond to averages for each fly pair (N=15 pairs). **J** Partner rotational speed during the transition to head interactions. Head interactions are driven by the partner male turning back to the courter during pC2l activation (increase in partner rotation). Solid lines correspond to mean and shaded area s.e.m. across all transitions (control: N=0 transitions, activation: N=4 transitions from 4 pairs). **K–N** Same as B–E, but for activation of pC1 neurons. Dots in L and N correspond to averages for each fly pair (N=7 pairs). **O** pC1 represents a broad set of neurons in the central brain of both females and males and consists of multiple subtypes expressing sex-determination genes doublesex (Dsx) and fruitless (Fru). A specific subset of neurons known as P1 expresses FruM proteins and is present only in males. P1a is a subtype of P1 neurons that drives courtship and aggression in males. **P–S** Same as B–E but for activation of P1a neurons. Dots in Q and S correspond to each fly pair (N = 15 pairs). **T** Same as **J**, but for P1a activation (control: N=4 transitions from 4 pairs, activation: N=28 transitions from 10 pairs). **U–X** Same as B–E but for activation of pC1x (pC1SS2) neurons. Dots in V and X correspond to each fly pair (N = 10 pairs). **Y** Putative circuit for song-induced feedback behaviors in male fruit flies. Distinct pathways drive chaining and head interactions, respectively, the former through the vPN1 neurons and a non-P1a pC1 subtype such as pC1x, and the latter through the pC2l neurons providing excitatory inputs to P1a. p-values are computed using two-sided Wilcoxon signed rank tests.

Male-specific vPN1 neurons are necessary for song-induced male chaining [52]. Chaining is only observed in groups of males and results in a chain of several males orienting head-to-tail [51, 53]. We tested whether the vPN1 neurons also induce head interactions between males. Activation of vPN1 in male pairs failed to induce head interactions and instead strongly increased tail interactions (Fig. 6B–E). Next, we tested the role of the pC2l neurons, which detect pulse song and induce acceleration and singing in solitary males [31, 43]. Unlike vPN1, optogenetic activation of pC2l neurons induced head interactions in male pairs (Fig. 6F–I), driven by the partner turning back to the courting male, as in wild types (Fig. 6J). Thus, distinct auditory pathways drive specific responses to song in males that lead to different modes of male-male interactions, with vPN1 driving tail interactions (Fig. 6B–E, [52]), and pC2l driving head interactions (Fig. 6F–J). In females, the pC2l neurons have been shown to induce slowing down [31]. Thus, the sexually dimorphic behavioral feedback to courtship song (males turning back and females slowing down) is mediated by the shared song detector neurons, pC2l [81]. The pC2l neurons drive song in males via the descending pIP10 neurons [82]. Recent studies suggest that pC2l neurons also connect to P1a neurons, a male-specific subset of the pC1 neurons, which accumulates social cues and encodes a persistent social arousal state [43, 83]. Another pC1 subset, called pC1x, is necessary for song-induced male-male aggression [45]. To test which of the pC1 subsets, P1a or pC1x, drive head interactions and courtship in male pairs, we activated the pC1 neurons or their subsets P1a and pC1x. Optogenetic activation of all pC1 neurons in pairs of males (Fig. 6K), each deafened to remove effects from endogenous song, promoted both tail and head interactions (Fig. 6L–M). Activation of the pC1x subset induced more tail interactions (head: 10 *±* 6% vs. tail: 90±6%, p=2e-3, Fig. 6U–X) and P1a activation (Fig. 6O—P) induced more head interactions (head: 68 *±* 24% vs. tail: 32 *±* 24%, p=0.01, Fig. 6Q–S), suggesting a specific role of P1a neurons in inducing head interactions in male pairs. Overall, the optogenetic activation data point to the existence of two pathways that link hearing song in male partners to specific feedback behaviors: First, one in which song is detected in vPN1 neurons to drive chaining, possibly via the pC1x neurons. And second, song detected by the pC2l neurons increases the male’s arousal and drives turning and singing towards other males via the P1a subset (Fig. 6Y).

## Discussion

Here, we have shown how the interplay between external behavioral feedback and internal sensori-motor processing shapes flexible social behavior by comparing courtship-like interactions directed at male and female partners. We found that the dynamics of courtship interactions change with the sex of the partner such that courting males were often facing the head of another male close up whereas often facing the tail of females (Fig. 1). This resulted in differences in the sequences of male- and female-directed courtship song with male partners receiving more sine song and females more pulses (Fig. 2). We then showed that courters use a fixed set of three sensorimotor rules for sequencing male- and female-directed song (Fig. 3). The differences in song sequences are driven by sex-specific partner feedback, which in turn drives sex-specific rule use by courters (Fig. 4). Sex-specific partner feedback is mediated by distinct neural pathways that link hearing song (Fig. 5) with behavior (Fig. 6). Together, these findings show that differential social feed-back by male and female partners can elicit differential actions through the same sensorimotor processing rules, leading to flexible context-dependent behavioral patterns.

We (Fig. 3) and others [13] have shown that *Drosophila* males pattern their courtship song using three sensorimotor rules, but how these rules are represented in the brain is unclear. Previous studies in non-human primates suggest that rule encoding in the brain manifests as correlated activity between sensory and action-related regions [84, 85]. Transitions between rules can alter these correlations through changes in sensory-motor cortical connections or through shared inputs from rule-encoding neurons to sensory and motor areas [84]. In rodents as well as human and non-human primates, neurons in decision-making regions such as the prefrontal cortex, orbitofrontal cortex (OFC), and striatum encode rule identity via changes in neuronal firing rates [5, 86–90]. Our findings on differential rule use in courting males according to partner behavior in *Drosophila* lay the groundwork for identifying the neural circuits that encode contexts and select rules in an experimentally accessible model organism with a known connectome [91].

In *Drosophila* males, the context-dependent rule use (Fig. 3) is driven not by the partner’s sexual identity but by their feedback (Fig. 4). Rule selection is therefore likely mediated by ascending and visual neurons that encode the locomotor state and the distance of the male courter from his interaction partner (Fig. 3M, S6B). Self-motion signals could be conveyed via central re-afferences [92] or from peripheral motor centers and proprioceptors [93–95]. High-order visual features that encode the distance and movement of the partner are processed in the lobular columnar neurons [96–98]. This context information is then likely integrated in the superior medial protocerebrum, likely involving subsets of pC1 and pC2 neurons (Fig. 6F–T) [13, 31, 43, 99]. Contributions of other integrative centers, such as the mushroom bodies or the central complex, cannot be ruled out [100]. All these circuits link sensory cues to motor outputs through descending neurons that target pre-motor centers in the ventral nerve cord [95, 101–103]. This is supported by recent studies showing that song patterns are shaped by differential recruitment of at least three types of descending neurons: pIP10, pMP2, and a descending neuron that is part of a disinhibitory circuit downstream of P1a [13, 43, 63].

The behavioral rules that specify how sensory cues are transformed into song can generate robust male- and female-directed song despite large differences in interaction dynamics, because the rules are sensitive to some, and invariant to other sensory cues: Song patterns are not shaped by the male position at the head or tail of the partner, but by differences in the partner’s speed and distance (Fig. 3, 4, S6). This implies that the male’s sensory experience during courtship can be split into two subspaces: an output-potent subspace (e.g., speed, distance) that drives singing, and an output-null subspace (e.g., angular position) that does not [104, 105]. This division enables males to generalize their singing to novel contexts, such as head interactions with other males. The concept of the output-null space originates in motor neuroscience, where it describes how neuronal activity in the primary motor cortex of non-human primates influences muscle activity during the execution, but not the preparation, of movement [104]. Preparatory neuronal activity resides in the output-null subspace, allowing planning without affecting execution. Similarly, in male singing, positional changes around the female [62] do not alter the song pattern because male position lies in the output-null subspace. This enables males to reposition to the female’s rear in preparation for copulation without disrupting their ongoing song pattern.

In our assays, a defining feature of male-male courtship-like behavior in *Drosophila* is the head interaction, initiated when the male partner turns back toward the courter. Both courtship song perception and heightened arousal are required to induce the partner’s turning (Fig. 5). Information about song in the partner is relayed via distinct auditory pathways with opposing effects: vPN1 drives tail chasing (Fig. 6C–E) and chaining [52], while pC2l drives head interactions (Fig. 6G– I). Our findings indicate that vPN1 and pC2l effect distinct behavioral responses via connections to different subpopulations of pC1 neurons: Effects of activation of the pC1x subset resemble those of vPN1 activation (Fig. 6D–E and W–X), suggesting a functional connection between both neurons, although this remains to be tested. By contrast, pC2l is thought to be connected to the P1a subset [6, 99], and indeed activation of both cell types elicits head interactions (Fig. 6G– H, Q–R). The P1a neurons control social arousal [78] and are part of an integrator network that might slowly accumulate song inputs from pC2l (Fig. 5M–N). Once the male’s arousal level is sufficiently high, the song-induced activity in P1a neurons surpasses inhibitory signals from male pheromonal cues [36, 106, 107], leading to disinhibition of the partner male’s courtship circuitry and male-directed singing. Once established, head interactions are likely stabilized internally by recurrent circuitry downstream of P1a [108] but likely also through a positive feedback loop formed by P1a driving song in one male, which activates P1a and drives singing in the other male. Further experimental validation is required to confirm the proposed neural circuit and song integration mechanism underlying male-male head interactions.

Overall, our findings demonstrate that sex-specific song patterning in *Drosophila* males emerges from context-dependent application of sensorimotor rules driven by sexually-dimorphic partner feedback. For example, female receptivity induced by male song slows her movement, triggering frontal circling where the male engages the *chasing* rule that elevates pulse song. On the other hand, male arousal induced by courtship song triggers a head encounter where the *close* rule is engaged, dominated by sine song (Fig. 7). The sexually-dimorphic song responses are likely mediated through sexually dimorphic song detection neurons in the pC2l and pC1 clusters (P1a in males and pC1a/c in females [31, 73, 81]). Although learning based adaptations to sensori-motor processing cannot be excluded, plasticity likely plays only a minor role under our conditions, since we used sex naïve flies. Instead, males flexibly recombine existing rules—exhibiting compositionality—a resource-efficient strategy for generating diverse behavioral sequences from a limited repertoire without the need for learning new actions [109–111]. While compositionality has been demonstrated in biological and artificial neural networks for cognitive decision-making tasks [109–112], our study extends it to natural behaviors. More broadly, our findings highlight how flexible, context-dependent behaviors can emerge from a static rule set via feedback-driven modulation, proposing a mechanism for flexibility of innate behaviors [2, 7–10, 113, 114]. Remapping existing rules to novel behavioral outputs likely also underpins the evolutionary diversification of behavioral repertoires [115, 116].

**Figure 7:**
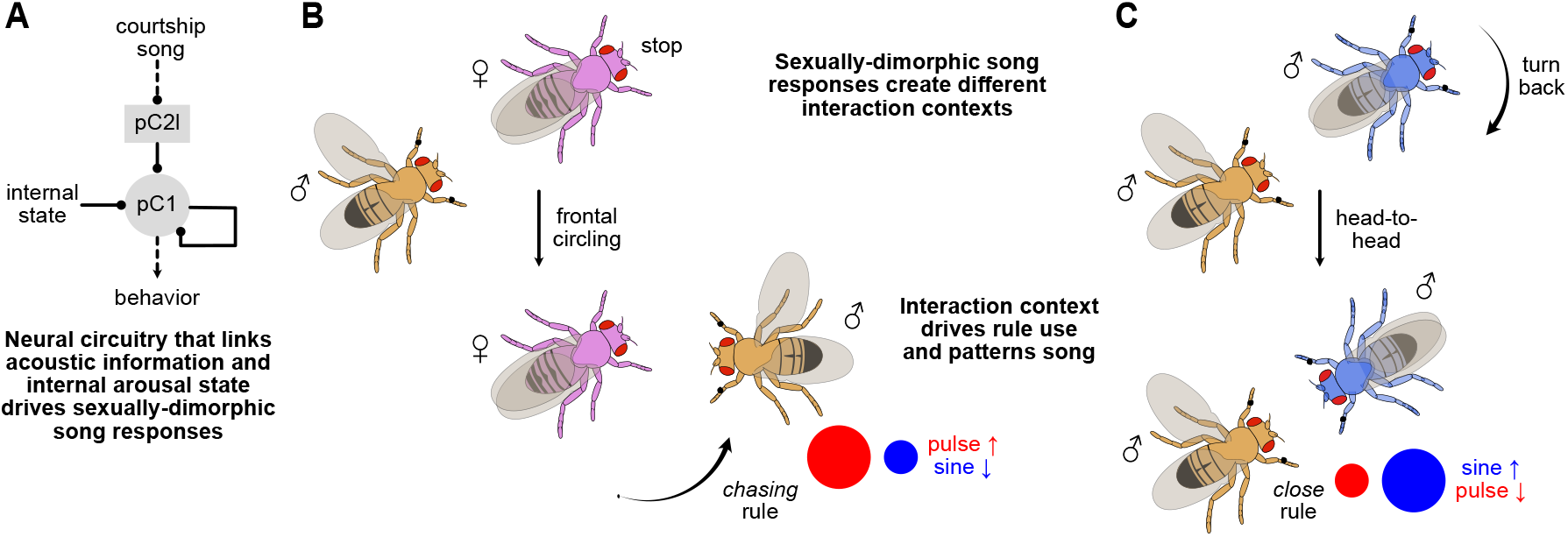
Sexually dimorphic behavioral responses to courtship song drive sex-specific rule use. **A** Courtship song is processed by the sexually dimorphic pC2l neurons [31] and pC1 neurons [52, 117]. Sex-shared pulse song detector neurons pC2l drive sexually dimorphic behavioral responses: Slowing down or stopping in females and turning back in males. **B** Song from a courting male induces sexual receptivity in the female partner and slows her, prompting the male to circle to her front. The courter predominantly uses the chasing rule to sing in this context. **C** On the other hand, song may induce sexual arousal in a male partner via pC2l and P1a neurons. He turns back and sings to the courter, establishing head interactions between males. The courter predominantly uses the close rule to sing in this context.

## Acknowledgments

We thank Frank Kötting and Stephan Löwe from the ENI workshop for their assistance in designing the behavioral chambers, Kimia Alizadeh for laboratory assistance, Lena Lindner for data acquisition, Gesa Hoffmann, Jan Schöning, Christine Gündner, Rüdiger Ludwig, Matthias Weyl, and Christiane Becker for technical and administrative support. We thank David Anderson, Vivek Jayaraman, André Fiala, Peter Andolfatto, Martin Göpfert, Janelia flylight, and Bloomington Stock Center for providing flies. We thank all members of the Clemens lab, as well as Daniela Vallentin, for feedback on the manuscript. This work was funded via an Emmy Noether Grant (Project number 329518246) and an ERC Starting Grant (Grant agreement No. 851210) to JC. SM and MN were funded by the IMPRS Neurosciences program of the University of Goettingen.

## Contributions

1. Conceptualization - SR, JC
2. Animals and behavioral experiments - SR, AP, SM, MN
3. Modeling - SR
4. Data analysis - SR, AP, SM, MN
5. First draft - SR, JC
6. Feedback on draft - AP, SM, MN

## Methods

### Experimental animals

Flies were raised on a standard cornmeal-agar medium at 25°C with 60% humidity and a 12-hour light/dark cycle. Within six hours of eclosion, virgin flies were separated by sex and housed in groups of 10–15 flies. For the experiments, we used flies that were between three to seven days old. For optogenetics, flies were housed on retinal food after eclosion. The retinal food was prepared by supplementing standard fly food with 400μM all-trans-retinal (Sigma-Aldrich R2500) dissolved in 99% ethanol. To prevent optogenetic activation, light exposure was minimized by wrapping the vials in aluminium foil.

**Table 1.**
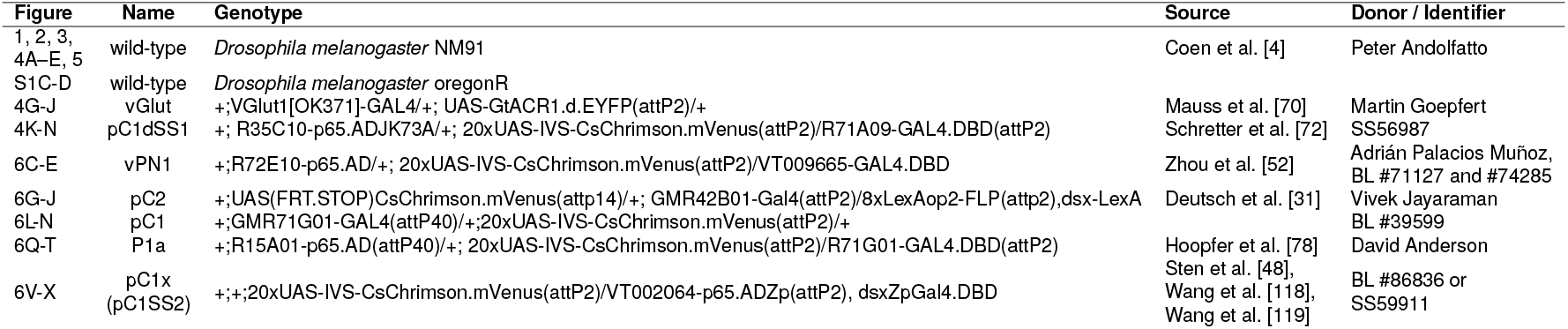
Fly lines used in this study.

### Behavioral assays

The behavior chamber was circular with a 44 mm diameter and 1.9 mm height unless otherwise specified. The large diameter allowed a wide range of fly behaviors and interactions whereas the narrow height prevented the flies from lifting off. The chamber and the lid were made of transparent acrylic to allow video recording. The lid was movable and contained a small opening for loading the flies. The chamber’s walls were sloped to prevent the flies from climbing to the ceiling. The floor of the chamber was tiled with 16 microphones (Knowles NR-23158) embedded in a custom PCB (Modified from Arthur et al. [55]). The acoustic signals were amplified using a custom-build amplifier (Modified from Arthur et al. [55]) and digitized using a data acquisition card (National Instruments PCIe-6343) at a sampling rate of 10 KHz. The microphones were covered with two layers of thin nylon mesh (50 mm diameter) for the flies to walk on. Fly behavior was recorded with a USB camera (FLIR flea3 FL3-U3-13Y3M-C, 100 frames per second (fps), 912 × 920 pixels) equipped with a 35 mm f1.4 objective (Thorlabs MVL35M1) mounted above the chamber. The chamber was illuminated using a circular array of white LEDs placed between the camera and the chamber. For optogenetic experiments, the chamber was illuminated with blue LEDs (470 nm, *1*.*5 µW* /*mm*^*2*^) whose wavelengths do not interfere with red and green wavelengths used for optogenetic manipulations. Song was played back using a loudspeaker (Hi-Vi Research^®^ Loudspeaker Model F6) placed on one side of the chamber.

Before the first experiment of each day, the chamber was prepared in two steps. First, the chamber and its lid were cleaned with ethanol, rinsed with water, and allowed to dry. The lid was then coated with Sigmacote (Sigma-Aldrich) to prevent flies from walking on the chamber ceiling. Second, to promote interactions between flies during the experiment, the chamber was perfumed with male and female pheromones by loading it with ∼5 male and ∼5 female flies for 5 minutes. For the experiments, flies were loaded individually into the chamber using an aspirator. Since the flies are most active at dawn, we conducted all experiments within 3 hours of the behavioral incubator lights switching on. Recordings were terminated when copulation occurred or after 15 minutes unless otherwise specified. Synchronized recordings of audio, video, and delivery of optogenetic stimuli were controlled using custom software (https://janclemenslab.org/etho).

We took several steps to promote courtship behavior in male-male pairs: First, we did not add a resource such as food or a decapitated or freeze-killed female, which typically drives aggression [44, 46, 48]. Second, since socially isolated flies are more aggressive, we group-housed males before the experiments [120]. Third, we perfumed the behavioral chamber with male and female pheromones by placing 5 males and 5 females into the chamber before each set of experiments [4]. Although perfuming may reduce the impact of volatile sex pheromones during courtship, interactions will still be influenced by the detection of non-volatile sex pheromones on the partner’s cuticle [121–123]. Lastly, we did not use genetic manipulations that interfered with the males’ ability to discriminate sex, since we wanted to study naturalistic male-directed singing. Instead, we used NM91, a strain of Drosophila melanogaster with a high motivation to court females [4].

### Optogenetic experiments

For the optogenetic experiments, flies were collected and placed on retinal food immediately after eclosion. For neural activation using csChrimson [71] or inactivation using GtACR1 [70], the chamber was illuminated using four red (625 nm) or green (525 nm) LEDs, respectively. A 500 nm short-pass filter (Edmund Optics, 500 nm 50 mm diameter, OD 4.0 Shortpass Filter) was attached to the camera’s objective to prevent the optogenetic stimulation from interfering with video recording.

#### Immobilization experiments

Partners were immobilized by using GtACR1 to inhibit all glutamatergic neurons, which include the motorneurons in the fly. To promote courtship towards the immobilized partners, we activated the P1a in the courter flies using csChrimson prior to the immobilization. The pair was initially exposed to red light for four minutes to induce a persistent arousal state in the courter by activating P1a neurons. Then, the pair was exposed to green light, to immobilize the male or female partners by inhibiting their motorneurons.

#### PC1d activation experiments

Wildtype NM91 males were paired with female flies expressing CsChrimson in pC1d neurons. Each experiment lasted 14 minutes, with the red light being OFF during the first 7 minutes (control) and ON during the next seven minutes (experiment).

#### pC2l and vPN1 activation

Two male flies that expressed uAS-csChrimson in vPN1 [52] (Fig. 6B–E) or pC2l [31] (Fig. 6F–J) neurons were paired for 30 minutes. The flies interacted for the first 15 minutes of the trial under normal light (control). Red light (27 *µW* /*mm*^*2*^) was turned ON for the next 15 minutes (experiment) which activated the vPN1 or pC2l neurons in the flies. Pilot experiments with pC2 activation in our large behavioral chamber (diameter 42 mm) showed that the flies frequently sang to the walls during activation. To promote interactions, we subsequently used a smaller chamber with a diameter of 16 mm and height of 3.25 mm.

#### pC1, pC1x and P1a activation

Two deaf male flies that expressed uAS-csChrimson in pC1 [52] (Fig. 6K–N), pC1x [48, 118, 119] or P1a [78] (Fig. 6P–X) were paired for 30 minutes. The flies interacted for the first 15 minutes of the trial under normal light (control). Red light (27 *µW* /*mm*^*2*^) was turned ON for the next 15 minutes which activated the pC1, pC1x, or P1a neurons (experiment). The flies were deafened by cutting their arista under cold anesthesia at least 12 hours before the experiment.

#### Auditory manipulations

For experiments in Fig. 5A–D, we paired a mute (wing-cut) male with an intact male (both NM91). For experiments in Fig. 5E–H, a deaf male was paired with an intact male (both NM91). For experiments in Fig. 5I–L, we used two mute (wing-cut) males (NM91). The flies were paired for 10 minutes, of which they interacted for the first 5 minutes in silence control. After 5 minutes, an artificial pulse song with a pulse width of 4 ms and an inter-pulse interval (IPI) of 36 ms was played back continuously for the next 5 minutes (playback experiment). For song playback, we used the protocol by Inagaki et al. [124]. We played back ten sound amplitudes at increasing volumes with each amplitude played back for ∼30 s. Manipulations (cutting the wings to mute males, cutting the arista to deafen males) were performed at least 12 hours before the start of the experiment under cold anesthesia.

#### Sexual satiation assays

To investigate the effect of sexual satiation, we used a satiation assay (Fig. 5O–Q). For these experiments, we paired a virgin mature male with a satiated male, both of which were 3 to 7 days old. The males were satiated by placing them in the presence of four females for up to 12 hours before the experiment. This allowed the males to copulated multiple times before the experiment and induced sexual satiation. To discriminate between the virgin and satiated males, the virgin mature males were marked with a white dot on their thorax using acrylic paint under ice anesthesia 12 hours before the experiment. To control for the ice treatment, the males to be satiated were also placed under ice anesthesia before placing them with the females.

### Annotation and analysis of the song

Acoustic signals were annotated using Deep Audio Segmenter (DAS, [56]), a deep learning-based tool for annotating multi-channel audio recordings. The acoustic recordings were segmented into four types – pulse song, sine song, agonistic song or wing flicks and noise (silence). To automatically annotate the recordings, the deep learning network was trained on a set of manually annotated recordings. The trained network was used to automatically annotate the remaining recordings which were then manually proofread.

We computed signal-level and pattern-level characteristics of the acoustic signals. To characterize pulse song, we extracted individual pulse waveforms from the song recording with a duration of 35 ms centered around each pulse center. The pulse carrier frequency was computed by performing a fast Fourier transform and computing the central frequency in the frequency spectrum with a cutoff at 1000Hz [41]. The pulse width was calculated based on the signal envelope, which was estimated using the Hilbert transform, and which was convolved with a Gaussian window of width 1.5 ms. The pulse width is then given by the duration the envelope remains above half of its peak amplitude. Pulses with a low signal-to-noise ratio or with multiple peaks were excluded from the analysis of pulse width. The pulse signals whose difference in maximum to mean envelope value is below a threshold of 0.08 was considered noisy and those pulses where the envelope crosses the threshold of 0.5 more than once was considered a multi-peak pulse. The pulse interpulse interval (IPI) was computed as the interval between to subsequent pulses. IPIs longer than 100 ms belong to different bouts and were excluded. The carrier frequency of sine song was based on the spectrogram of individual sine waveforms obtained by dividing each sine bout into segments of 256 samples (∼ 25 ms) and calculating the modal central frequency across segments.

We discriminated the two pulse types *P*_*f ast*_ and *P*_*slow*_ (Fig. S3C–D) using unsupervised clustering of the pulse waveforms. We normalized [125], embedded the pulse waveforms into a UMAP, and clustered them using the H-DBSCAN algorithm [59, 125, 126]. We considered the two biggest clusters produced by H-DBSCAN, and labelled all pulses within each cluster based on the shape of the waveforms in each of the clusters. The data in the remaining clusters were labeled by training a support vector classifier (SVC [127]) using the labels obtained from the two biggest clusters.

We also computed the amount, the number of onsets, and the duration for pulse trains, sine songs, and all song (combining pulse and sine) in overlapping windows of 1 minute (50% overlap). All song bouts were defined as a sequence of pulse trains and sine song that were interleaved by less than 100 ms of silence. The amount was given by the fraction of the window occupied by pulse, sine, or all song. The number of onsets is given by the number of transitions into pulse, sine or any song. The bout duration is computed as the ratio of the amount and the number of onsets in each window. For each quantity, the average over all windows for a given fly was computed.

The bout order is given by one plus the number of song-mode transitions within a song bout. A bout order of 1 corresponds to bouts with only pulse or only sine. A bout with a single transition between pulse and sine has bout order two, and so on.

### Pose tracking and processing

The position of six body parts (head, neck, thorax, abdomen, tips of the left and right wing) was tracked across frames for each fly using Deepposekit [128]. For most of the analyses, the tracking data was down-sampled from the original frame rate of 100 Hz to 30 Hz. The tracks were transformed into a set of 10 metrics describing the locomotion and relative positioning of the two flies in the chamber. The fly velocity in the x- and y-direction were computed as the rate of change of the fly thorax position along the respective directions. The forward velocity was (cFV/pFV; c: courter, p: partner) was computed as the component of the velocity vector in the direction of the fly’s body axis (defined by the line connecting head and abdomen). Lateral velocity is the velocity component perpendicular to the body axis. Due to the left-right symmetry, we consider only the magnitude of lateral velocity—the lateral speed (cLS/pLS). The angle of each fly is computed as the angle spanned by the body axis and the image x-axis (positive is counter-clockwise). The rotational velocity of the fly is computed as the rate of change of that angle. Again, due to left-right symmetry, only the magnitude of the rotational velocity, the rotational speed (cRS/pRS), is considered. The distance between two flies (dis) was computed as the Euclidean distance between their thorax positions. The relative angle of one fly with respect to another (c*θ* and p*θ*) is computed as the angle between the body axis of the partner and the line connecting the thoraces of the two flies. The relative orientation (*ϕ*) is computed as the difference between the body angles of the two flies. These metrics function as input features to both the HMM-GLM model that predicts the male song patterning as well as the unsupervised social maps.

For detecting unilateral and bilateral wing extensions (Fig.1C, S3A), the angle of each wing was computed as the angle between the line joining the head and thorax and the line joining the thorax and the corresponding wing tip. A unilateral wing extension occurs when one of the wing angles exceeds a specific threshold. For each trial, this threshold is set at the wing angle that maximizes the number of frames correctly identified as song frames compared to human annotations. A bilateral wing occurs when both the left and right wing angles exceed a threshold of 30 °.

#### Quantifying courter position during interactions

We used a polar histogram to quantify the position of the courter around the partner. The distance was measured in terms of the partner fly length. The radial position axis was limited to three fly lengths and binned in 15 equal intervals. Since the side (left or right) on which the courter is placed is not relevant, we used absolute values of relative courter angle *cθ* for the angular position. The angular position was thus limited from 0 ° to 180 ° and divided into 36 bins of 5 ° each. *f θ*=0 ° specified the head and *f θ*=180 ° specified the tail (abdomen) of the partner. The quadrant spanning *cθ*=0 ° to *cθ*=90 ° was considered as the *head* quadrant (the courter was placed near the head of the partner) and the quadrant spanning *cθ*=90 ° to *cθ*=180 ° was considered as the *tail* quadrant (the courter was placed near the tail of the partner). To compute the probability of being placed in a quadrant during the interaction, that is P(quadrant|interaction), we ignored the bouts in which the courter was situated in a given quadrant for less than 0.5 s to remove short transients and possible effects of noise. Similarly, if the time elapsed between the end of a bout and the start of the next bout in the same quadrant was less than 0.5 s, they were considered to be the same bout. After this, the P(quadrant|interaction) was computed as the ratio of frames in which the courter was placed in a given quadrant and the frames in which the courter was interacting with the partner.

#### Transitions from tail to head interactions

We defined a tail interactions as instances when the courter was within 8 mm from the partner fly (*dis < 8 mm*), the partner was within a field of view of 60 ° (*pθ*<60 °), and the courter was positioned in the tail quadrant of the partner fly (90 °<*cθ*<180 °). Head interactions were identified when *dis < 8 mm, pθ*<60 °, and the courter was positioned in the head quadrant (*cθ*<90 °). We identified transitions from tail to head interactions when the courter, after interacting in the tail quadrant for at least two seconds, moved into the head quadrant and remained there for at least two seconds.

We further quantified the probability of song in time windows of different durations leading to transitions to head interactions. For each transition, we considered a window of *L* seconds before the transition and computed the probability of singing pulse, sine, or either in this window across all such transitions. We compared this with the probabilities of song in the windows that did not lead to a transition. For this, we considered 100 random windows of length *L* per experiment where the endpoint of the window was at least one minute away from a transition. To ensure that the effect was not linked to the difference in overall interaction between the flies during these windows, we ignored windows in which the interaction ratio was less than 25%.

### Social maps

We constructed social maps by jointly embedding the dynamical tracking features from both interaction partners into a low-dimensional space. For a experiment of duration *T* and a time history of *K* = *15* samples (0.5 ms) for *M* = *10* features, we created a design matrix of size (*T, Kx M*) by flattening the time history of all features into a vector. We then fitted a nonlinear dimensionality reduction algorithm, UMAP [59], using 10% of uniformly temporally spaced samples from all experiments of both male-female and male-male interactions to obtain a two-dimensional embedding space. We then embedded the remaining data from all the trials into this low-dimensional space. Thus, for each trial we generate an embedding of size *T × 2* where *T* is the number of samples in a trial. With the data pooled from all the trials, the two-dimensional embedding space functions as a state space that captures the social interactions with similar interactions clustered together and different interactions embedded away from each other. A density function of the samples in this state space characterizes the probability of visiting that region of the state space. To create a continuous state space representation, we perform a kernel density estimation over the samples in the low dimensional representation. Depending on the frequency of certain interactions, this state space consists of regions with higher probability (stereotypic interactions that occur often) and regions with lower probability (for example, rarely occurring interactions or transitions between two stereotypic interactions). Thus a probability-aware segmentation of the interaction state space would generate distinct regions belonging to distinct interaction contexts between the animals. To achieve this, we perform a watershed segmentation over the inverted density estimate of the UMAP-embedded samples [58]. We confirmed by manual inspection that the different watershed clusters correspond to stereotypic interaction contexts.

### Maps of behavioral responses to song

To analyze the responses of male and female partners to courtship song, we generated behavioral maps based only on the partners’ egocentric tracking features. A frequency spectrogram was computed from each pose coordinate data (normalized using fly length to account for variations in fly size) using a continuous Morelet wavelet transform which returned the signal power at 25 dyadically spaced frequencies ranging from 1Hz to 25Hz. Thus for each fly, we obtain a time series of dimension *T*× *24*× *25* where *T* is the trial duration. This time series is then flattened into a dimension of *T* ×*600* which is then embedded into a two-dimensional space using manifold embedding technique UMAP [59]. As before, we trained the UMAP embedding algorithm using 10% of data (uniformly sampled in time) from all the flies and used the learned UMAP to embed the rest of the data, performed kernel density estimation and partitioned the state-space into stereotypical behaviors using the watershed transform.

### HMM-GLM modeling

We used an HMM-GLM to model the sensorimotor rules that map the feedback cues to the male’s choice of song mode (pulse, sine, or silence) at each time point following Calhoun et al. [13]. An HMM-GLM model is a combination of a hidden Markov model (HMM) and a generalized linear model (GLM). An HMM identifies hidden states from observations (here mode of the male courtship song). In a vanilla HMM, at each time step, the model is in one of the hidden states and has a fixed probability of transitioning to another state or staying in the same state. Similarly, for each state, there is a fixed probability of observing an outcome (a song mode). Although the hidden states identified by the HMM may reveal the internal states of the animal associated with the observed behavior, it does not take into account the effect of external stimuli on the behavior. To account for this, an HMM-GLM associates with each hidden state a GLM that maps the history of sensory cues at each time point to the song mode probability. As surrogate sensory cues, we used the 10 metrics extracted from pose-tracking data. These metrics describe the locomotion and relative positioning of the two flies and thus function as proxies for the feedback cues that a courter male would receive when patterning the song. For each of the 10 metrics, we used a stimulus history of 4s (120 samples at 30 Hz) as input to the model at each time point. Thus at each time point, the model receives a feature vector of length 1200.

#### Hidden Markov model (HMM)

A Markov process or a Markov chain **z** is a stochastic process that describes a sequence of states of a system in which the probability of being in a particular state at any given time *t* depends only on the state occupied in the immediate previous time *t* − *1*.

In a hidden Markov model (HMM), the stochastic process underlying the sequence of states is not directly observable (hidden), but can only be observed via another stochastic process **y** whose outcome depends on the hidden Markov process **z**. Let {*z*_*1*_, *z*_*2*_, … *z*_*T*_} represent a sequence of hidden states of **z**, and {*y*_*1*_, *y*_*2*_, … *y*_*T*_} represent the corresponding observations from **y**. In our study, the hidden states represent a distinct sensorimotor rule that the male used to pattern his song at each time point, whereas the observation refers to a discrete song mode (pulse, sine or silence) that the male produced at each time point.

An HMM is characterized by the number of discrete hidden states *K*, the number of distinct observations *M* (here the number of song modes), an initial probability distribution *π*∈ ℛ^*K*^ (where *K* is the number of hidden states), whose each element gives the probability to be in a state at *t* = *1*, with *π*_*i*_ = *P*(*z*_*1*_ = *i*), a state transition matrix *α ∈* ℛ^*K×K*^ whose elements *α*_*i,j*_ == *P*(*z*_*t*_ = *j*|*z*_*t−1*_ = *i*) gives the probability of transitioning from one hidden state *i* at time *t − 1* to another hidden state *j* at time *t*, and an observation matrix *η ∈* ℛ^*M×K*^ whose each element *η*_*k,j*_ = *P*(*y*_*t*_ = *k*| *z*_*t*_ = *j*) gives the probability of observing an observation *k* when the system is in state *j*. Therefore an HMM can be completely specified by its parameters as

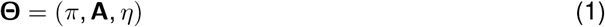

#### Generalized linear model (GLM)

To incorporate the effect of external sensory inputs on the male’s song patterning behavior, we replaced the fixed probability observation matrix *η* ∈ ℛ ^*M×K*^ by a generalized linear model (GLM) which maps the history of feedback cues at each time point **s**_*t*_ *−* _*N*_, …, **s**_*t*_ to a song mode probability *P*(*y*_*t*_ = *i* | **s**_*t* − *N*_, …, **s**_*t*_). Similar to the Calhoun et al. [13], we used a multinomial GLM which outputs probabilities of three types of song: pulse, sine, and silence at each time point. For simplicity, we considered fast and slow pulses [41] as a single pulse mode. The feedback history **s**_*t*_ − _*N*_, …, **s**_*t*_ is convolved with a set of *D* basis functions. Each basis function is a vector **b**_*j*_ ∈ ℛ ^*N*^ of dimension equal to the cue history *N*. We used raised cosine functions [129] that broaden with delay to capture the fact that the effect of cues is often strongest close to the behavioral response (small delays). The basis transformation serves two purposes: 1) It smoothens the feedback cues and thus reduces the effect of noise on the model, and 2) it reduces the dimensionality of model inputs from *10 × N* to *10 × D* (usually *D ≪ N*). Thus the transformed feature vector **x** = {**s**_*t−N*_, …, **s**_*t*_} *·* {**b**_*j*_ ∈ ℛ, *j ∈* {*1*, … *D*}} has a length of *10 ×D*. This feature vector is z-scored and is augmented with a ‘1’ to incorporate a bias, thus yielding a vector of length (*10* × *D*) + *1* as input to the GLM. The transformed input **x** ∈^(*10 ×D*)+*1*^ is passed through linear filters **w**_*i*_ ∈^ℛ (*10 ×D*)+*1*^, *i* ∈ {*1, 2, 3*} corresponding to three song modes, and a softmax function which maps the vector of filtered feedback cues to a normalized probability measure of each song type at each time point *t*.

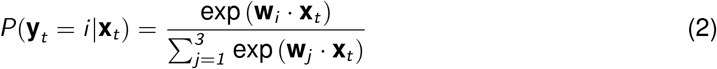

The filters for one song type (here silence) are set to zero such that probabilities of all song types sum to 1. While a previous study that used HMM-GLM [13], used a penalty on the difference of filter coefficients to ensure smoothness, we used basis functions as filters (see also Coen et al. [4]) to reduce the dimensionality of the inputs and ensure smoothness. An L2 regularization was applied to the filter coefficients.

#### HMM-GLM

The HMM-GLM we used here associates a GLM parameterized by its filter coefficients **W**_*k*_ = {**w**_*i*_ *∈* ℛ ^(*10 ×D*)+*1*^, *i ∈* {*1* … *M*}}, with each hidden state *k* of the HMM. Similar to a vanilla HMM, the transition from one hidden state to another follows a fixed probability distribution *α ∈* ℛ ^*K×K*^ and an initial state distribution *π ∈* ℛ ^*K*^. Thus, the parameters of HMM-GLM can be written as

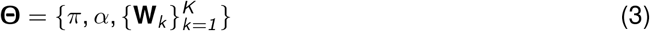

The optimal parameters of the model are found by maximizing the log-likelihood of the observed song sequences given the model parameters and the feedback cues.

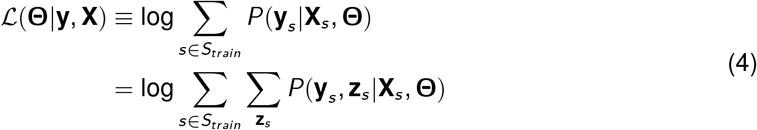

where *S*_*train*_ ⊂ *S* represents the sessions (trials) whose data (feedback cues and song) is used for learning the HMM-GLM parameters. The above equation requires summing across all possible state combinations **z**_*s*_ (paths through the hidden Markov chain) for each trial which is of exponential complexity. The forward-backward algorithm proposed by [130] solves it recursively thus reducing the complexity to linear in *T*. The HMM-GLM was trained using the Expectation-Maximization (EM) algorithm proposed by [131].

#### Forward-Backward algorithm

The forward-backward algorithm was proposed by [130] and provides an efficient way to compute the log-likelihood function in Eq. 4. It involves computing the “forward” and “backward” probabilities. The forward probabilities *a*_*i,t*_ give the probability of all observations up to time *t* and the system is in state *i* at time *t* (trial index *s* omitted for clarity).

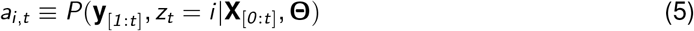

The backward probabilities *b*_*i,t*_ give the probability of all future observations up to time *T*, if the state at time *t* is *i*.

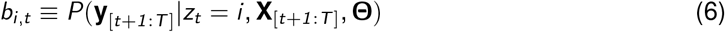

The forward and backward probabilities can be computed recursively by induction [132], thus reducing the complexity to linear in *T*.

#### Expectation-Maximization (EM)

**E-step:** The E-step involves computing the posterior distribution of the sequence of hidden states given the observations and the model parameters, *P*(**z** | **y, X, Θ**). From the forward and backward probabilities, we can compute the marginal distribution of the states at every time step and the marginal distribution of the state transitions as

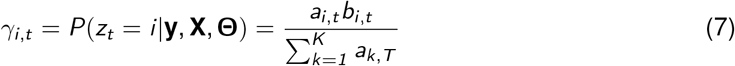

and

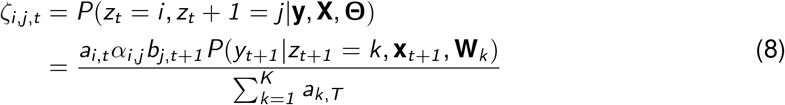

**M-step:** The M-step involves computing the parameters **Θ** that maximize the expected loglikelihood given the observations and the posterior state distribution computed in E-step.

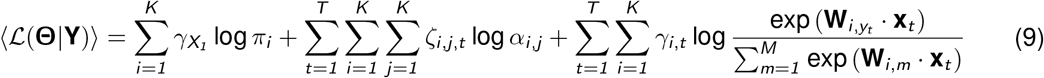

The E- and M-steps are iterated until convergence is reached. Since the algorithm is susceptible to getting trapped in local minima giving trivial solutions, we fitted the model multiple times with different initialization of model parameters.

#### Chance model

A chance model gives the probability of observing a song mode *m*, at the current time point *t*, as

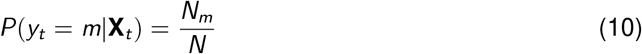

where *N*_*m*_ is the number of frames for which song mode *m* is observed, and *N* is the total number of frames in the trial. Thus a chance model always outputs the average probability of observing a song mode and ignores the feedback cues.

#### Testing

Once the HMM-GLM parameters were learned, the model performance was validated by computing the log-likelihood of observations (song sequences) on held-out trials.

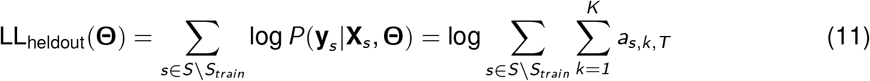

The normalized log-likelihood gives the improvement in prediction of song mode compared to a chance model and is obtained as

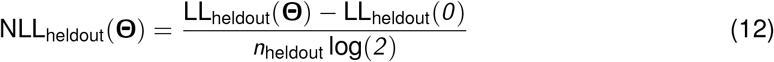

where *n*_heldout_ is the number of heldout samples and LL_heldout_(*0*) is the log-likelihood obtained from a chance model.

The fitted HMM-GLM can also be used to infer the sequence of hidden states for the complete trial given the observations and feedback cues using the E-step (Eq. 7) or the Viterbi algorithm [133]. Here the inferred states refer to the distinct sensorimotor rule that the male used to pattern his song at each time point. Once the hidden state sequence is inferred, it can also be used to predict the probability of each song mode at each time point by applying the filters corresponding to the inferred state to the feedback cues at that time point.

#### Sex-agnostic and sex-specific models

To identify whether the males use the same set of sensorimotor rules to pattern their courtship song towards female and male partners, we fitted three models: 1) a model fitted to female-directed song, **Θ**_*f*_, 2) a model fitted to male-directed song, **Θ**_*m*_, and 3) a model fitted to both male- and female-directed song, **Θ**_*a*_. Further, we validated the performance of sex-specific models on the same sex (held-out trials) and opposite sex and the sex-agnostic models on each sex separately (on held-out trials). For example, the performance of the three models in explaining the female-directed song was given by LL_*f*_ (**Θ**_*f*_), LL_*f*_ (**Θ**_*m*_), and LL_*f*_ (**Θ**_*a*_). If these three likelihoods are similar, a model with sex-specific rules does not explain the observed song patterns better than a model with sex-agnostic rules.

#### Feature importance

To evaluate which feedback cues are relevant for song patterning, we performed a feature importance analysis. This was done by randomly shuffling each of the ten features keeping the other features and song patterning data unchanged and then evaluating the model performance on the shuffled data. Feature importance was computed as the log ratio between the log-likelihood of the model on the shuffled data and the original data. The lower the ratio, the higher the importance of the given feature.

#### Effect of courter position and orientation on model predictions

To evaluate the effect of courter male position and orientation around the partner on HMM-GLM predictions of song mode, we computed the difference in predicted probabilities of pulse and sine song by the model using original and simulated positions and orientations of courter male. For each original sample, we create 180 simulated samples by keeping all the features except either courter relative angle (*cθ*, for courter position) or partner relative angle (*pθ*, for courter orientation to partner) and relative orientation (*ϕ*) unchanged. We then sweep *cθ* (or *pθ*) from 0 ° to 180 ° (1 ° increments). Correspondingly *ϕ* is set to *180* ° −*cθ* (when sweeping *f θ*) or *180* ° −*pθ* (when sweeping *tθ*). For each simulated sample, the model-predicted probability of pulse and sine song was compared to the predicted probabilities for the original sample. The difference in predicted probabilities is then visualized as a function of the original courter position around the partner.

#### UMAP visualization of filtered feedback cues

We collected the filtered feedback features by applying the learned GLM filters based on the inferred state (rule), *z*_*t*_, and the observation (song mode) *y*_*t*_ to the basis transformed inputs **x**_*t*_ at each time point *t*. A GLM sums up these filtered feedback cues and passes them through a non-linearity (logistic function) to output a probability for singing a song mode. Therefore, these filtered features represent a decision space that the model inferred to be relevant for song patterning with boundaries in this space illustrating a change in decision between one song mode versus the other. To visualize such a decision space, we reduced the dimensionality of the filtered feedback cues from (*10* × *D*) + *1* to *2* using UMAP [59]. This analysis showed that similar regions in the decision space map to the same song outputs during both female- and male-directed courtship (Fig. S6G).

#### Inferring states from optogenetic experiments

We used the HMM-GLM models fitted to the wild-type song patterning data (**Θ**_*a*_) to infer the rules used by males during optogenetic experiments.

### Statistical analyses

All statistical tests were performed using a two-sided Mann-Whitney U (unpaired data) or two-sided Wilcoxon signed-rank (paired) test. A significance level of 0.05 was used.

### Softwares and algorithms

**Table 2.** Software and algorithms used.

| Resource | Identifier |
| --- | --- |
| DeepPoseKit | https://github.com/jgraving/DeepPoseKit [128] |
| DeepAudioSegmenter (DAS) | https://github.com/janclemenslab/das [56] |
| GLM utilities | https://github.com/janclemenslab/glm_utils |
| scikit-learn | https://scikit-learn.org [134] |
| HMM-GLM | https://github.com/lindermanlab/ssm/ |
| umap-learn | https://github.com/lmcinnes/umap [59] |
| FFTKDE | https://github.com/tommyod/KDEpy |
| scikit-image | https://scikit-image.org/ [135] |
| python 3.7 | https://python.org |
| Scipy 1.7 | https://scipy.org [136] |
| NumPy 1.21 | https://numpy.org/ |
| pandas 1.3 | https://pandas.pydata.org/ |
| seaborn | https://seaborn.pydata.org/ [137] |
| matplotlib | https://matplotlib.org/ [138] |
| statannotations | https://github.com/trevismd/statannotations [139] |
| Affinity Designer | https://affinity.serif.com/designer/ |

## Supplementary Figures

**Figure S1:**
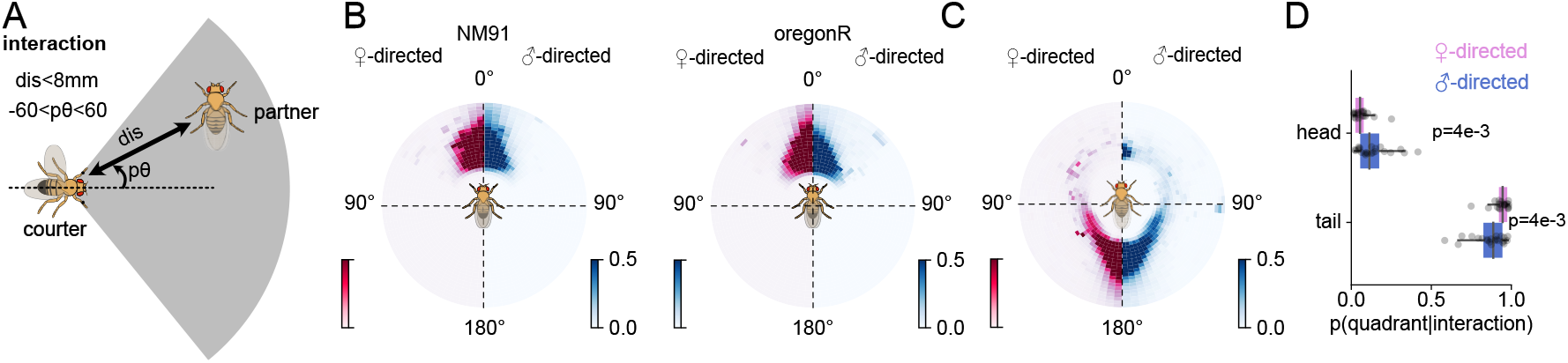
Fly orientation during interactions. **A** Definition of interaction, courter (or focal) and partner (or target) flies. An interaction occurs when the flies are within 8 mm of each other and one fly (the partner or target) is within the field of view (± 60°) of the other fly (courter). When both flies are within a field of view of less than ± 60° of each other, the fly that was initially courter remained so. **B** Position of the female and male partner fly with respect to an NM91 (left) or OregonR (right) courter male. The courter fly is always oriented towards the partner during interactions. **C** Position of an OregonR courter male around a female (left, magenta) or a male (right, blue) OregonR partner during interactions. 0° and 180° represent the partner’s head and tail, respectively. **D** Ratio of time spent by the OregonR courter male near the head and tail of the OregonR partner fly during interactions. p-values were computed using a two-sided Mann-Whitney U test.

**Figure S2:**
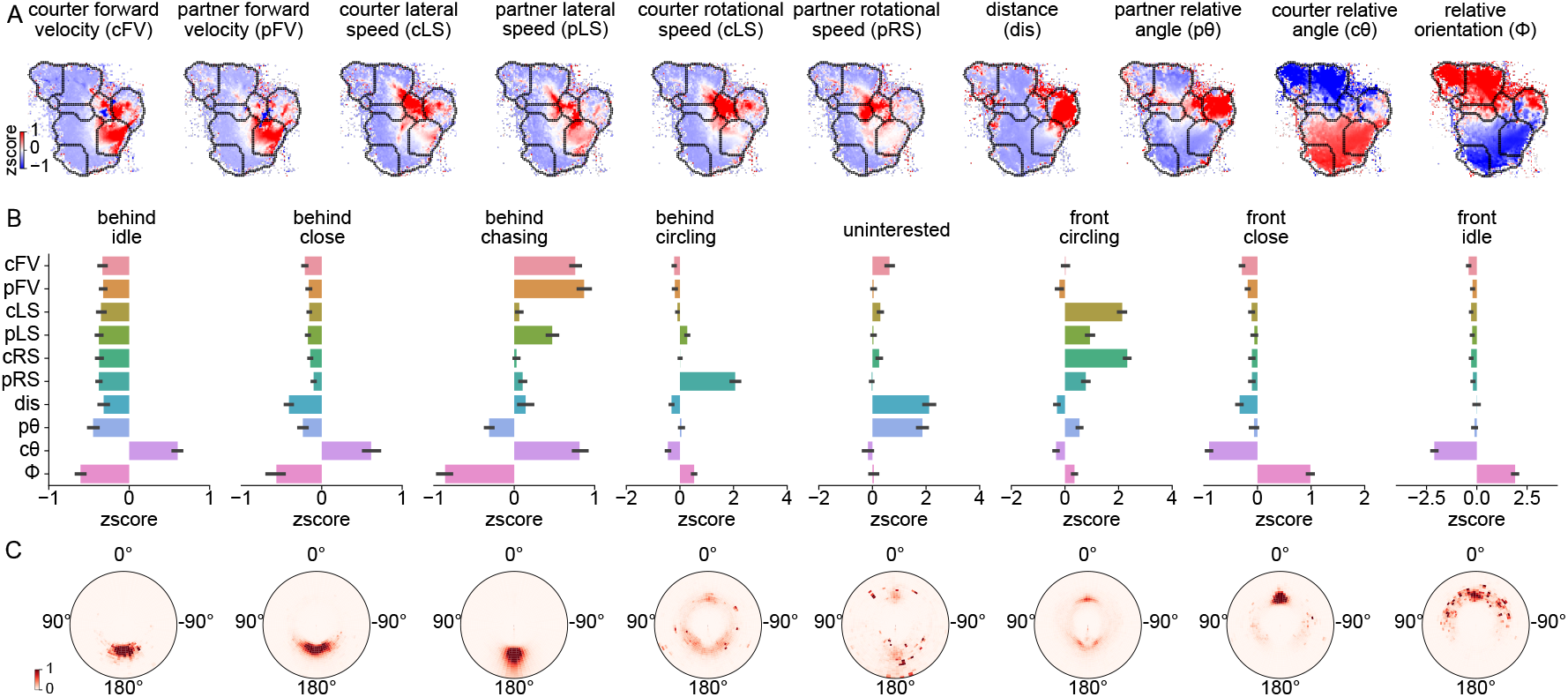
Unsupervised quantification of male- and female-directed interactions. **A** Behavioral features for different locations in the social map. Color shows the z-scored behavioral feature values for each pixel in the map. **B** Behavioral features in each of the eight social modes. Bars and error bars show mean ±standard deviation across fly pairs. Features were z-scored. **C** Position of the courter around the partner during each of the eight social modes. In the first four modes (behind idle, behind close, behind chasing, behind circling), the courter is mostly placed behind the partner. For the last three modes (front circling, front close, and front idle), the courter is mostly placed near the partner’s head. The radial axis is normalized to the target length and is limited to three fly lengths.

**Figure S3:**
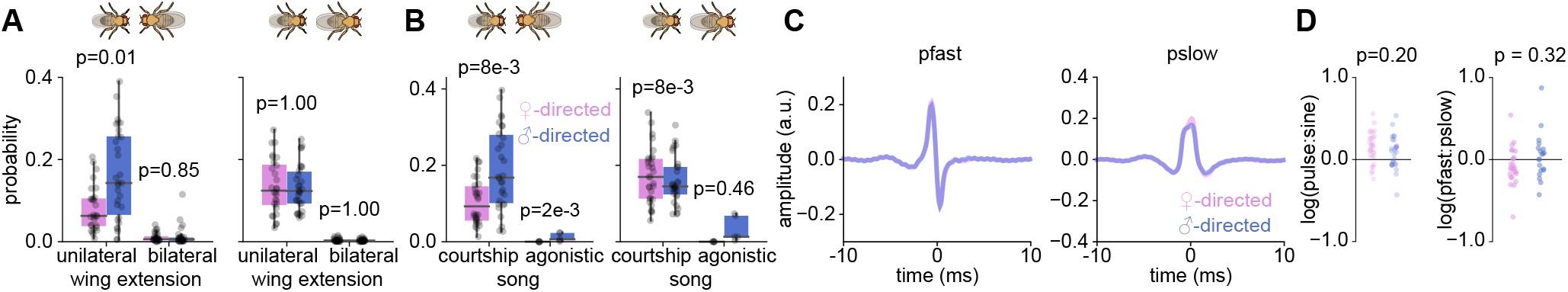
Female and male-directed singing were courtship-like with little difference in short-term song characteristics. **A** Unilateral and bilateral wing extensions during tail (left) and head interactions (right) with a female (magenta) and male (blue) partner. **B** The amount of courtship and agonistic song during tail (left) and head interactions (right). **C** The average waveform of fast (Pfast: N=23288 for female-directed and 24913 for male-directed) and slow pulses (Pslow: N=21329 for female-directed and 21374 for male-directed) sung towards female and male partners. The pulse shape does not change with partner sex. **D** Pulse-to-sine (left) and fast-to-slow pulse (right) ratio during female- and male-directed interactions. p-values are computed using a two-sided Mann-Whitney U test. Dots in A, B, and D correspond to the average value for each fly pair (N=5 pairs each of male-female and male-male agonistic song. For everything else, N=30 pairs each of male-female and male-male).

**Figure S4:**
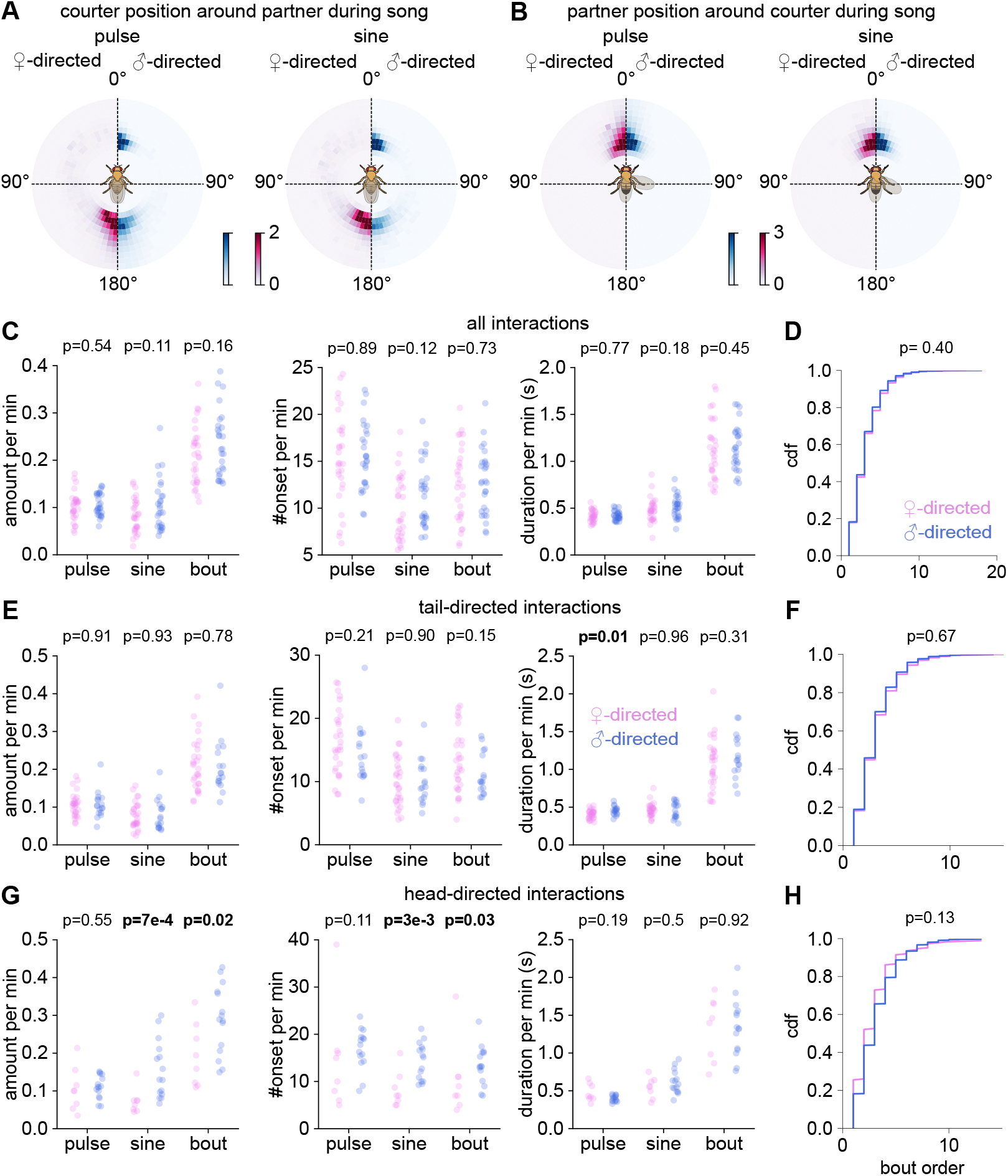
Song patterns differ most strongly during head interactions. **A** Courter position around the partner during singing pulse (left) and sine (right) song. The courter is situated mostly near the tail when singing to a female, but also near the head when singing to a male. The courter is situated at a broader range of distances and angular positions when interacting near the tail of a female than the tail of a male. **B** Partner position around courter during pulse (left) and sine (right) song. Both songs are produced when the partner is within a narrow field of view of the courter (narrower during male-directed singing). **C** Amount, number of onsets, and duration of pulse, sine, song bouts per minute during male-(blue) and female-directed (magenta) interactions. The overall difference between female- and male-directed song patterning statistics is small. Dots correspond to the average value of song metrics for each fly pair (N=30 pairs each of male-female and male-male). **D** Cumulative density functions of the bout order for male (blue) and female-directed (magenta) song. Bout order corresponds to the number of transitions between song modes (pulse, sine) within a song bout. **E–F** Same as C and D restricted to tail interactions. Dots correspond to the average value of song metrics for each fly pair with tail-directed interactions (N=30 pairs each of male-female and male-male). **G–H** Same as C and D restricted to head interactions. Dots correspond to the average value of song metrics for each fly pair with head-directed interactions (N=8 pairs of male-female and 15 pairs of male-male) All p-values are computed using a two-sided Mann-Whitney U test.

**Figure S5:**
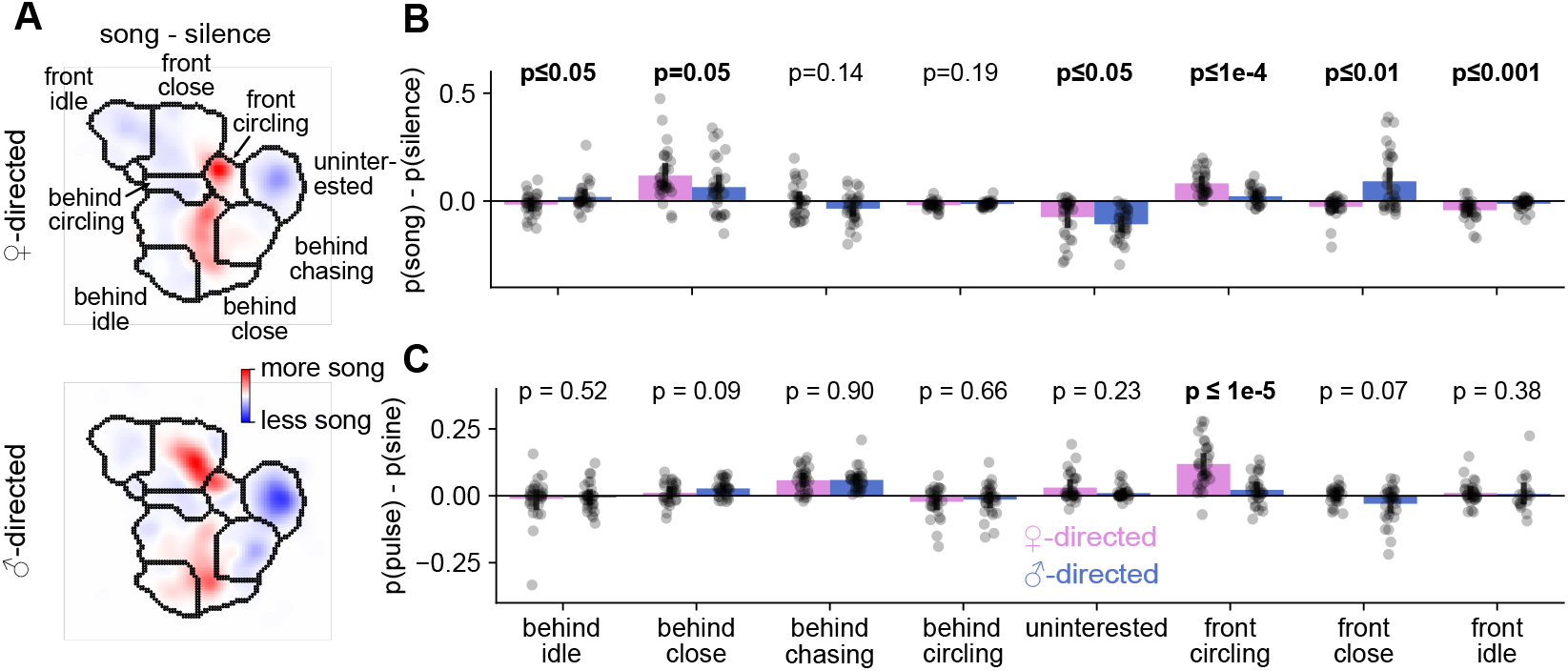
Males sing in different social modes towards two partners. **A** Difference between the interaction state spaces conditioned to song (pulse or sine) and silence during female-(top) and male-directed (bottom) interactions showing context-specific differences in singing based on partner sex. When interacting with females, males produce more song during the close, chasing, and frontal circling modes. When interacting with males, males sing more during the behind and idle, behind and close, and frontal close modes. **B, C** Difference between the probabilities of singing and staying silent (B, data from A) and pulse and sine (L, data from Fig. 2F–G) in different social modes. Dots correspond to average values for each fly pair (N=30 pairs each of male-female and male-male). Bar heights and error bars correspond to the average ±std across fly pairs. All p-values are computed using a two-sided Mann-Whitney-U test.

**Figure S6:**
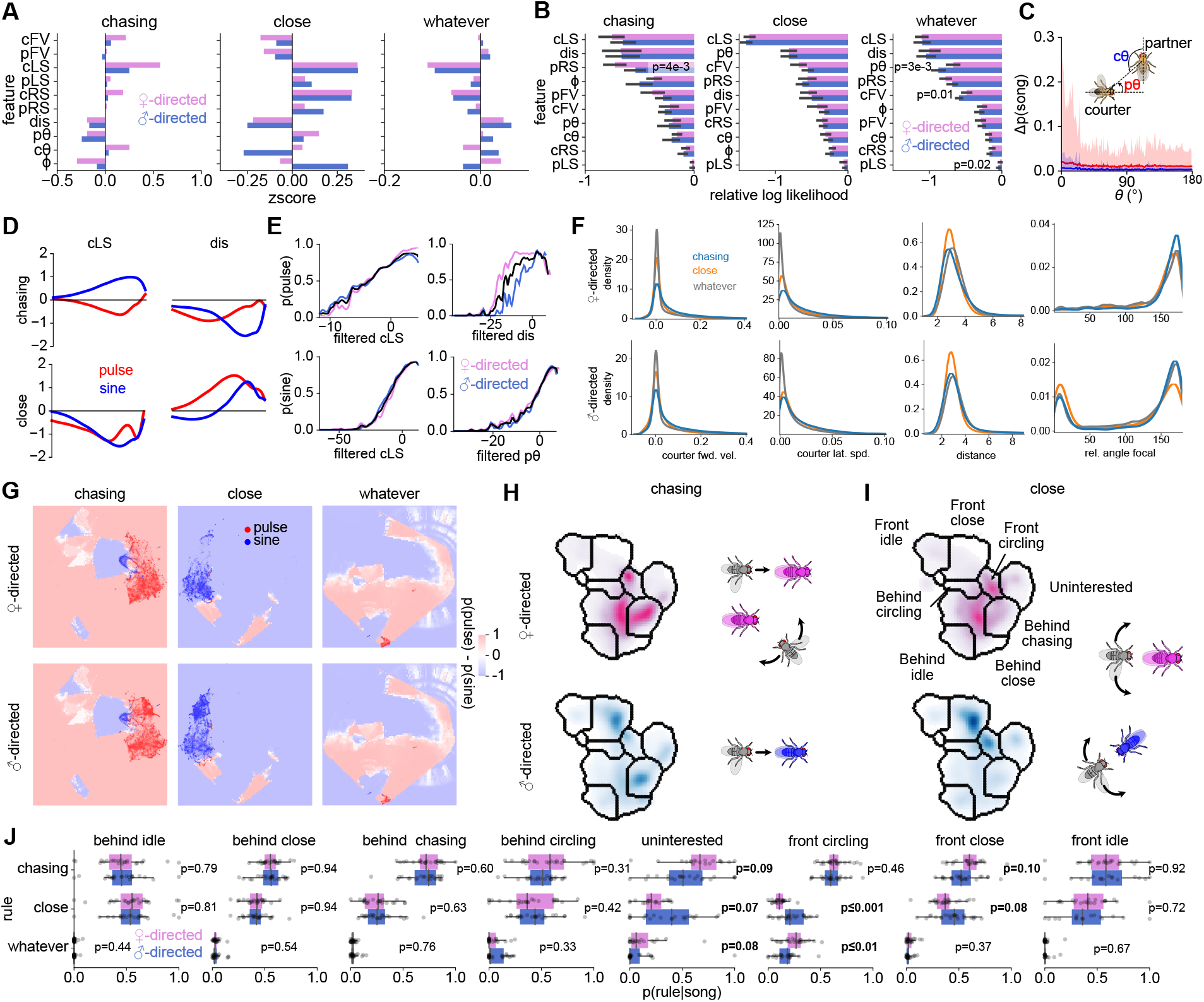
Modeling song patterning rules using HMM-GLM. **A** The mean of z-scored features during the chasing, close, and whatever rules. **B** Mean±std importance of behavioral features for each rule in the HMM-GLM, computed as the reduction (logarithmic scale) in model log-likelihood when the feature is randomly permuted. **C** Effect of courter position and courter orientation around the target on HMM-GLM predictions, computed as the difference in predicted probabilities of song by the model when changing the relative courter angle (*cθ*) between 0 (head-directed) to 180deg (tail-directed) or relative partner angle (*pΘ*) between 0 (focal oriented towards the target) to 180deg (focal oriented away from the target). The solid curves show the mean, and the shaded regions show the 95% CI. When changing *cθ*, the mean change is virtually zero and the 95% CI is less than 0.04, indicating that the male’s angular position around the target has a negligible impact on song patterns. **D** Filters for courter lateral speed (cLS) and distance (dis) for pulse (red) and sine (blue) during the chasing and close rule. **E** Probability of pulse (top) and sine (bottom) song as a function of filtered cLS and dis during the chasing and close rule. Curves are similar for female-(magenta) and male-directed (blue) singing, indicating that the mapping from behavioral features to song is independent of sex. Tuning curves for male and female directed pulse song are slightly offset for distance, due to differences in the distance distributions, with males tending to sing pulse song even at farther distances from females than from males (Fig. S4A–B). **F** Probability density of courter forward velocity (cFV), courter lateral speed (cLS), distance (dis), and relative courter angle (*cθ*) during the chasing, close, and whatever rules during female (top) and male-directed interactions (bottom). For both male and female targets, the courter male is fast and far during the chasing rule, close and slow during the close rule, and slow and far during the whatever rule. Additionally, towards male partners, the courter is predominantly placed near the head during close state. **G** Filtered stimuli space visualized using UMAP embedding along with decision boundaries learned by the HMM-GLM visualized in the UMAP space (see Methods). The background colors show the difference between pulse and sine probabilities inferred by the model in the stimulus space. **H, I** Social maps (left) and illustration of the interaction contexts (right) when the male uses the chasing (**H**) and close (**I**) rules during interactions with female (top row, magenta) and male (bottom row, blue) partners. **J** Rule use for each social mode. The rule use depends on partner sex only in the modes associated with head interactions. Dots correspond to average value for each fly pair (N=30 pairs each of male-female and male-male pairs). All p-values were computed using a two-sided Mann-Whitney U test.

**Figure S7:**
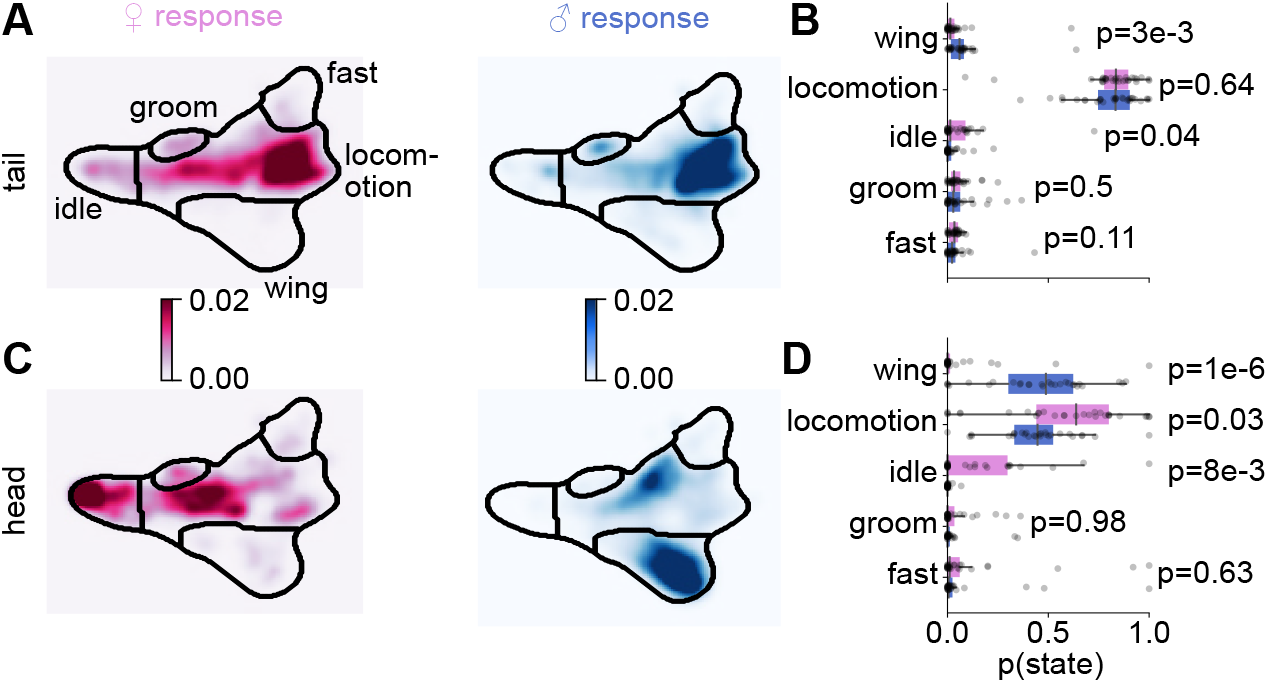
Partner’s behavioral feedback drives differential song patterning. **A–B** Probability of different behaviors of female (right) and male (left) partner while the courter male is singing in the tail quadrant. The higher duration of sine song towards female partners during tail interactions (Fig. 2E) may be attributed to the increased idleness of females during such interactions which allow the male to interact more closely near female tail (Fig. 1H). **C–D** Same as **A–B** but when the courter male is singing in the head quadrant. Differences in partner behavioral feedback are much more pronounced when the courter is in the head quadrant, driving differences in head-directed song patterning. Dots in **B, D** correspond to each fly pair (N=30 male-female and 30 male-male pairs). All p-values were obtained using a two-sided Mann-Whitney U test.

**Figure S8:**
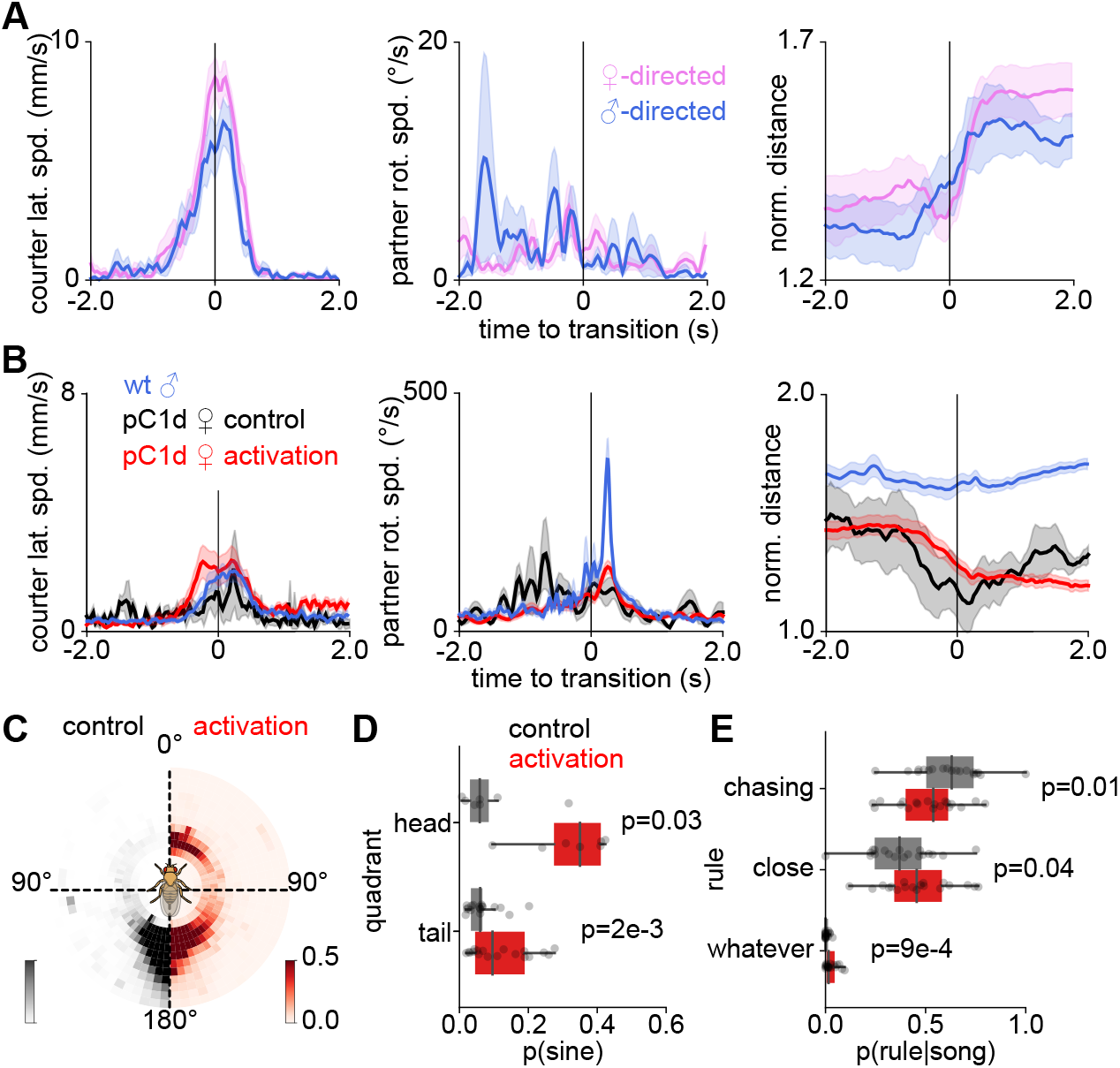
Manipulating partner’s behavioral feedback modulates rule use and song patterning. **A** Courter lateral speed (left), partner rotational speed (middle), and normalized distance (right) during transitions into head interactions with an immobilized female or male partner. Both female- and male-directed transitions occur when the focal fly circles to the front of the target and thus leads to an increased distance between flies. Lines and shaded areas correspond to mean ±s.e.m. (14 transitions from 10 male-female pairs and 16 transitions from 9 male-male pairs). Distance is normalized by the body length of the immobilized partner. **B** Courter lateral speed (left), partner rotational speed (middle), and distance to the partner (right) during transitions in head interactions with a female partner expressing csChrimson in aggression-inducing pC1d neurons. During activation (LED on), the partner turns back like a wild-type male partner reducing the distance between the flies. Lines and shaded areas correspond to mean ±s.e.m. (control: n=6 transitions from 5 pairs, and activation: n=57 transitions from 17 pairs). **C** Position of a male courter around a pC1d-Chrimson female during control (grey, LED off) and activation (red, LED on). **D** Fraction of sine song produced by male courter in the head and tail quadrants around a pC1d-Chrimson female during control (grey) and activation (red). **E** Rule use when singing to a pC1d female partner during control (LED on) and activation (LED off). Dots in D and E correspond to each fly pair that interacted in the given quadrant during both control and activation (Head: N=8, Tail: N=19). For panels D–E, p-values were obtained using Wilcoxon signed-rank tests.

